# The initiation of *de novo* protein folding on the ribosome

**DOI:** 10.64898/2026.07.24.740641

**Authors:** Ivana V. Bukvin, Julian O. Streit, Tomasz Włodarski, Charity R. Hornby, Sammy H.S. Chan, Anaïs M.E. Cassaignau, John Christodoulou

**Author notes:** These authors contributed equally to this work (Ivana V. Bukvin, Julian O. Streit).

## Abstract

How the earliest structure within the unfolded state is formed during biosynthesis on the ribosome and whether it has any consequences for downstream folding remain open questions. Here, we combine ^15^N paramagnetic relaxation enhancement NMR with all-atom molecular dynamics simulations to characterise the unfolded state of a folding-competent immunoglobulin-like domain on the ribosome at the cusp of folding initiation. We identify three structurally distinct sub-ensembles that differ in compaction and ribosome interactions. Non-native contacts, together with ribosome interactions, likely delay folding, yet their persistence alongside early native-like contacts within a sparsely populated compact sub-ensemble suggests they may also facilitate the formation of a co-translational folding nucleus, whose contacts overlap with those of the downstream intermediates. From these findings we infer a mechanistic model of *de novo* folding initiation during biosynthesis and, by linking the folding nucleus to downstream partially structured intermediates and the native state, provide a complete atomistic description of a co-translational folding pathway.

## Introduction

AlphaFold has revolutionised our capability to predict the final fold of many structured proteins in recent years^1^. However, the pathway by which a protein reaches its final fold in cells remains a fundamental open question^2^. Although elegant *in vitro* refolding experiments over the past seven decades provided insights into the pathways and energetics that govern protein refolding from a denatured to a native state^3–6^, *de novo* protein folding in the crowded cellular environment is increasingly being shown to differ significantly from refolding in dilute aqueous solutions, and the underlying mechanisms remain largely unexplored^7–11^.

In particular, the molecular determinants governing how the first structural elements emerge within the unfolded state during biosynthesis on the ribosome remain elusive^6,12^. Generally, unfolded states explore a rough energy landscape with many local minima, characterised by an ensemble of highly disordered dynamically interconverting conformations of similar energies, and readily perturbed by solvent and transient interactions^13^. Consequently, insights gained through refolding studies^14–30^ may not fully capture the conformational preferences and dynamics of the unfolded state during *de novo* protein folding^31^.

Most proteins can fold co-translationally, and some are obligate co-translational folders^9,32,33^. The ribosome can modulate the thermodynamics^34–40^ and (un)folding rates^36,41^ of nascent chains (NCs) through interactions^35,42–44^ and steric restriction^45–47^. In cells, every polypeptide chain is synthesised by the ribosome in a vectorial manner, and most initially explore, at least in part, disordered conformations during their biosynthesis^9,39^. We recently demonstrated how the ribosome destabilises these unfolded states^40^, at least for negatively charged domains^39^, which promotes the formation of downstream partially structured intermediates^34,48^. The structural basis for how folding is first initiated within the unfolded state itself however remains unknown.

We combine paramagnetic relaxation enhancement (PRE) NMR with all-atom molecular dynamics (MD) simulations to characterise the unfolded state of a folding-competent immunoglobulin-like domain, FLN5, on the ribosome at the cusp of folding initiation. We identify three structurally distinct sub-ensembles, comprising a dominant ribosome-anchored state, an expanded state, and a sparsely populated compact state, each shaped by a distinct interplay of ribosome interactions, non-native contacts and early native-like contacts within the nascent chain. The compact sub-ensemble, enriched in native-like contacts, likely seeds the formation of a co-translational folding (coTF) nucleus whose contacts overlap with those of the downstream partially structured intermediates. Although the role of non-native contacts in protein folding has been debated^25,26^, we find that both non-native contacts and ribosome interactions are present across all three sub-ensembles, likely modulating the coTF landscape in a way that delays premature folding while ultimately facilitating the formation of the coTF nucleus, thus participating in productive *de novo* folding on the ribosome.

## PRE NMR and MD reveal structures of the unfolded state at the cusp of folding on the ribosome

We investigated the unfolded state of a folding-competent immunoglobulin-like (Ig-like) domain, FLN5^34,35,42,45,49–51^, on the ribosome (Figure **1a**). For the ribosome-nascent chain complex (RNC), FLN5 is tethered to the ribosome peptidyl transferase centre (PTC) via a 31-amino acid linker (FLN5+31), comprising the subsequent FLN6 domain and SecM AE1 stalling sequence (Figure **1a**). While this NC snapshot has the entire FLN5 sequence emerged from the ribosomal exit tunnel^42^, it nevertheless significantly (44 ± 8%) populates an unfolded state^34^. ^15^N NMR data exclusively report on the unfolded state, as the more structured intermediates (totalling 46 ± 9%) and native states (10 ± 4%) remain invisible due to (i) their slower molecular tumbling^42^ and (ii) the very slow (k_ex_ < 1 s^−1^) exchange between the unfolded and the structured states^34^. PRE-NMR experiments of the FLN5 RNC (Extended Data Figures **1**-**3**) show multiple long-range PRE effects, suggesting the conformational ensemble is exploring transient long-range intramolecular contacts (Figures **1b**, **1c** and Extended Data Figures **1a**, **2**). By contrast, a folding-incompetent A3A3 variant, in which two clusters of aromatic residues in the core of the domain were mutated to alanines thus abolishing folding ability without significantly perturbing the unfolded state ^35,39^, predominantly explores short-range PRE effects (Figures **1b**, **1c**, Extended Data Figure **1a**)^39,52^.

**Figure 1.**
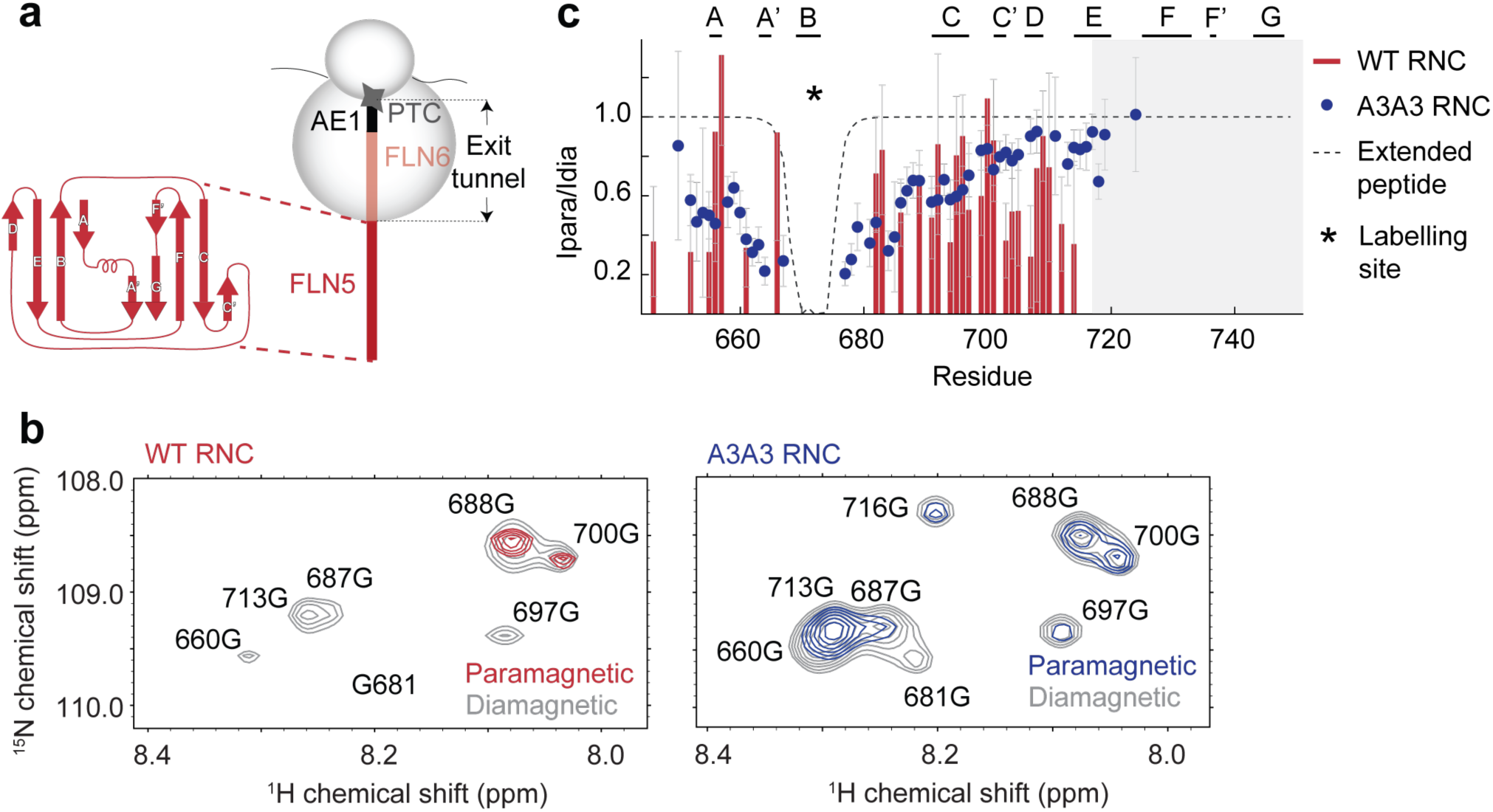
The unfolded state of the wild-type, folding-competent, FLN5 RNC explores long-range PREs compared to the A3A3, folding-incompetent, variant. **a**, Schematics of the constructs used for the PRE FLN5+31 experiments. The RNC is comprised of an N-terminal 6xHis-tag, the FLN5 domain, the subsequent FLN6 domain, and the SecM-AE1 stalling sequence^34,35,39^; the inset depicts the structure topology representation of the native FLN5 domain with secondary structural elements (PDB 6G4A^51^). b, Representative region of a ^1^H-^15^N SOFAST-HMQC NMR spectrum of the FLN5+31 C671 (left) and A3A3+31 C671 (right) RNCs with the paramagnetic and diamagnetic spectrum overlayed (see Extended Data Figure 2 for full spectra). NMR data were recorded at 800MHz, 283K. c, PRE-NMR intensity ratio profiles for FLN5+31 C671 (red) and A3A3+31 C671 (blue, see Extended Data Figure 1b-c for all labelling sites). Theoretical reference profiles corresponding to a fully extended polypeptide are also shown as dashed lines. Native FLN5 secondary structure elements (β-strands) are annotated at the top. The shaded region represents the C-terminal region of FLN5 that is broadened beyond detection due to ribosome interactions (via residues 728-747)^35^. The black asterisk indicates the position of the MTSL labelling site. Errors were derived from the standard deviation of the spectral noise.

Structural restraints from these experiments were used to reweight an ensemble obtained from all-atom MD simulations of the FLN5 unfolded state on the ribosome with explicit solvent, as previously described (Extended Data Figures **4a-b**, **4g-h**)^39^. The reweighted structural ensemble is in good agreement with the PRE-NMR data and is validated against experimental chemical shift data on the ribosome (Extended Data Figures **4a**, **4c**, **4d**).

## Folding competence is associated with enhanced entropic destabilisation of the unfolded state on the ribosome

We have previously described how the ribosome entropically destabilises the A3A3 folding-incompetent unfolded state primarily through increased solvation arising from NC conformational expansion^39^. To determine whether this extends to the folding-competent wild-type FLN5 unfolded state, we compared its structural ensemble to the isolated wild-type counterpart obtained using all-atom simulations as previously described for the A3A3 unfolded state in isolation (Extended Data Figures **4e-f**, **4i**, **5l-m**)^39^. This isolated ensemble was in good agreement with experimental NMR chemical shifts, residual dipolar couplings^51^ and the radius of hydration^51^ determined for the Δ6 FLN5 variant (C-terminal truncation; Extended Data Figures **5h-k**), which represents the closest approximation to a wild-type unfolded state in isolation.

Analysis of the thermodynamic parameters of the ensembles revealed that the wild-type FLN5 unfolded state is destabilised on the ribosome relative to in isolation, due to a reduction in both conformational entropy (+2 ± 1 kcal mol⁻¹, Extended Data Figures **6a–f**) and solvation entropy estimated from changes in solvent-accessible surface area (SASA +19 ± 7 kcal mol⁻¹; Extended Data Figures **6g**, **6i**). Overall, the unfolded state free energy is estimated to be increased by +16 ± 9 kcal mol⁻¹ on the ribosome relative to the isolated ensemble, after accounting for enthalpic solvation effects (Extended Data Figure **6h**). As previously observed for the A3A3 variant^39^, the ribosome entropically destabilises the unfolded state, though the effect is ∼80% larger than that estimated for the A3A3 unfolded state ensemble (+9 ± 6 kcal mol⁻¹), suggesting that the degree of entropic destabilisation of the unfolded state on the ribosome may be intrinsically linked to folding competence.

## The unfolded state on the ribosome explores three structurally distinct sub-ensembles

Comparing the obtained wild-type, folding-competent, FLN5 unfolded state ensemble to the A3A3 variant (Figure **2a**)^35,39^, allows us to identify the structural characteristics of the unfolded state specifically associated with folding competence. Overall, the two ensembles show similar levels of expansion relative to the isolated polypeptide (Extended Data Figures **4e-i**)^39^, with the A3A3 unfolded state being just ∼4% more expanded with an average radius of gyration (Rg) of 44.1 ± 1.8 Å compared to 42.1 ± 1.7 Å (mean ± s.d. of the distribution) for the wild-type FLN5 unfolded state on the ribosome (Extended Data Figure **5a**). Both ensembles predominantly sample random coil-like structures with a minor contribution of β-sheet secondary structure, which is somewhat more prevalent in the FLN5 ensemble (1.6 ± 0.2% and 1.2 ± 0.1% for the FLN5 and A3A3, respectively), with considerable variation in their distribution along the primary sequence (Extended Data Figure **5e**). The average intrachain contact probabilities, including both short- and long-range contacts, were comparable, 4.6 ± 0.2% for FLN5 and 4.4 ± 0.2% for A3A3. Despite these overall similarities, the distribution of transient native and non-native contacts differed substantially (Figure **2c**), suggesting that folding competence may also be associated with the specific pattern of contacts sampled within the unfolded state^53^.

**Figure 2.**
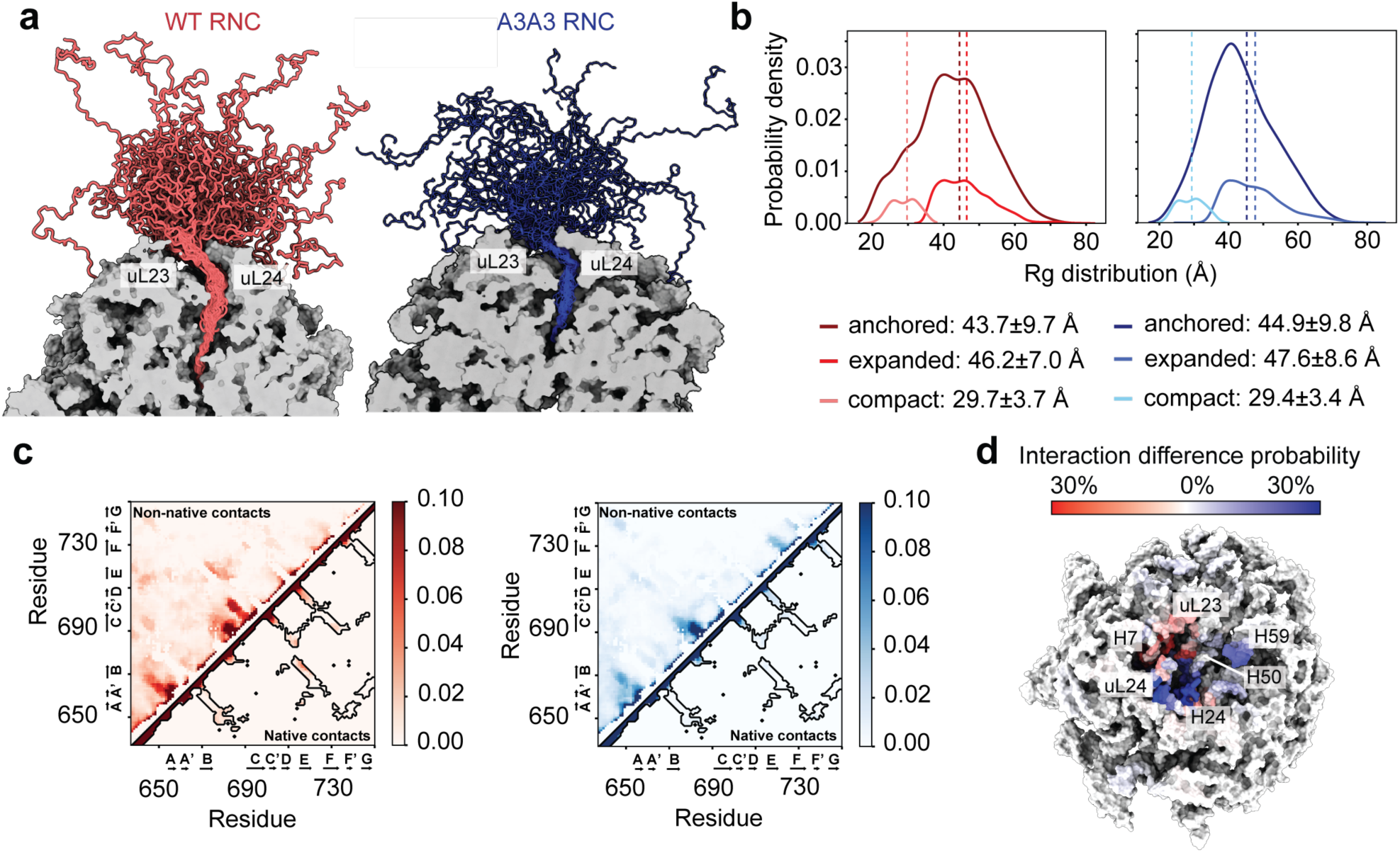
Comparison of the PRE reweighted structural ensembles of the wild-type FLN5 and A3A3 unfolded states on the ribosome. **a**, Representative ensembles (top 50 cluster centroids after reweighting with the PRE NMR data) of the wild-type FLN5 (left) and A3A3 (right) unfolded state on the ribosome. The ribosome cross-section is shown in grey and is sliced to the exit tunnel to enable visualisation of the linker within the tunnel. **b,** Radius of gyration probability distributions of the ribosome-anchored, expanded and compact structural sub-ensembles of the wild-type FLN5 (left; 77.1%, 13.2% and 9.7%, respectively) and A3A3 (right; 82.0%, 13.4% and 4.6%, respectively) unfolded states on the ribosome. **c,** Average contact map of the wild-type FLN5 (left) and A3A3 (right) unfolded state on the ribosome (zoomed in to a probability of 0.1). Contacts were defined as Cα-Cα distances within 10Å and recapitulated when using closest heavy atom definitions (cut-off 6 Å; Extended Data Figure 9b). Non-native contacts are mapped above the diagonal; native contacts are mapped below the diagonal. The black contours highlight the native contact map of the folded FLN5 domain. **d,** Difference in NC-ribosome contact probability mapped on the ribosome surface, highlighting regions contacted more by FLN5+31 (red) and A3A3+31 (blue). Cα atoms of the NC and ribosomal proteins and P, N3, and C4’ atoms of the rRNA, and a distance cut-off of 12.5 Å were used to define NC-ribosome contacts. rRNA helices and ribosomal proteins surrounding the exit tunnel are annotated.

We next explored the ribosome interaction probability of both the wild-type FLN5 and A3A3 unfolded state ensembles and found that 77% of the FLN5 and 82% of the A3A3 structures interact with the ribosome via a C-terminal binding segment (residues 728-747; Extended Data Figures **5b-c**)^35^, forming the most populated sub-ensemble in both FLN5 and A3A3 RNCs, termed ‘ribosome-anchored’ (Figures **2b**). Two distinct exit pathways have been previously identified for NCs within the ribosomal exit vestibule, mediated by interactions with the H6/H7 and H24/H50 regions of the ribosome^52^. Although our ribosome-NC PRE NMR data report exclusively on the emerged portion of the NC and do not resolve interactions within the exit tunnel (Extended Data Figure **1a-b**), the ribosome surface interaction profiles of both the wild-type and A3A3 unfolded state ensembles engage both regions of the ribosome surface, but with distinct preferences: the wild-type favouring the H6/H7 and the A3A3 the H24/H50 region of the ribosome surface (Figure **2d**, Extended Data Figure **5d**).

Within the remaining fraction of the ensembles, we identify two additional distinct sub-ensembles: a more expanded population, termed ‘expanded’, and a more compact population, termed ‘compact’ (Figure **2b**), with the two populations separated at an Rg of 36 Å and the compact sub-ensemble occupying a distinct low-energy region of the free energy landscape that is less accessible in the folding-incompetent A3A3 variant (Figure **2b** and Extended Data Figure **5f-g**). The separation of the two sub-ensembles by degree of compaction was further corroborated using the energy landscape visualisation method (ELViM; Extended Data Figures **7a-f**)^54^. Together, these distinct sub-ensembles form the basis for understanding early folding events, and we investigate their structural characteristics and role in *de novo* protein folding initiation on the ribosome.

## Non-native interactions and ribosome-binding can delay folding initiation in the most-populated sub-ensemble

To explore the influence of the observed ribosome binding on the initiation of FLN5 folding, we initially examined the structural features of the ribosome-anchored FLN5 unfolded state sub-ensemble (Figure **3a**) which was determined to comprise 77% of the wild-type FLN5 unfolded state ensemble on the ribosome, and exhibiting an average Rg of 43.7 ± 9.7 Å (Figure **2b**). It predominantly explores local non-native contacts, in particular between the loop connecting strands B and C, and strands C and C’, the loop connecting strands B and C, and strand C, and, to a lesser extent, between strands A and A’ at the N-terminus, as well as transient native contacts (0.6 ± 0.1%) between strands A and B, B and E, and C and E (throughout, strand nomenclature refers to sequence regions that form β-strands in the native FLN5 fold but remain largely unstructured within the unfolded state ensembles; Figure **3b**). These contacts form predominantly between polar and hydrophobic residues (Figure **3c** and Extended Data Figure **9a**). The β-sheet secondary structure propensity (1.5 ± 0.2%) aligns with the regions exploring non-native contacts, including strands A and E, which form transient native contacts (Figure **3d**). By comparison, the folding-incompetent A3A3 ribosome-anchored sub-ensemble displays a somewhat reduced β-sheet propensity (1.2 ± 0.1%), which is best defined in the loop region connecting strands B and C and in strand C (Extended Data Figure **8b**). It explores non-native contacts between strands A and A’, the loop connecting strands B and C, and strand C, as well as between strands F and F’, and F’ and G, but does not explore the non-native contacts between the loop connecting strands B and C, and strands C and C’ (Extended Data Figure **8a**).

**Figure 3.**
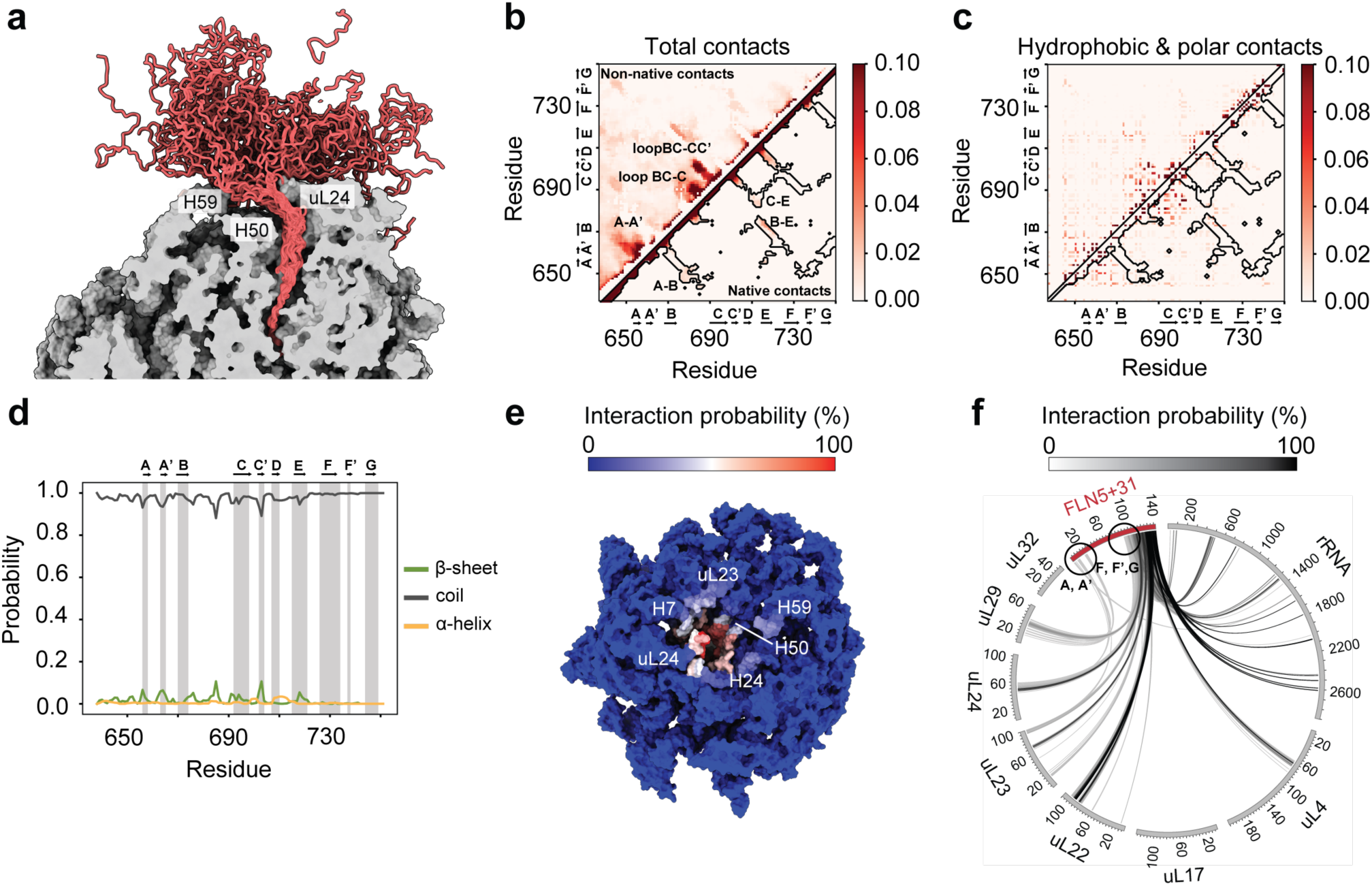
Structural preferences of the ribosome-anchored sub-ensemble of the FLN5 unfolded state on the ribosome. **a**, A representative sub-ensemble (top 50 cluster centroids after reweighting with the PRE NMR data) of the ribosome-anchored unfolded state of wild-type FLN5 on the ribosome. The ribosome cross-section is shown in grey and is sliced to enable visualisation of the FLN6 linker within the tunnel. Average contact maps of the ribosome-anchored sub-ensemble of the wild-type FLN5 unfolded state on the ribosome (zoomed in to a probability of 0.1) indicating total contacts **(b)** and contacts between hydrophobic and polar residues only **(c)**. Contacts were defined as Cα-Cα distances within 10Å and recapitulated when using closest heavy atom definitions (cut-off 6 Å; Extended Data Figure 9b). Non-native contacts are mapped above the diagonal; native contacts are mapped below the diagonal in **(b)**. The black contours highlight the native contact map of the folded FLN5 domain. **d,** Average secondary structure propensities in the ribosome-anchored sub-ensemble of the wild-type FLN5 unfolded state on the ribosome along the protein sequence determined using the DSSP algorithm. The vertical shaded areas highlight the regions of β-strands (annotated as strands A-G) in natively folded FLN5. **e,** Interactions of the wild-type FLN5 NC with the ribosome mapped on the ribosome surface in the vicinity of the exit tunnel^39^ as observed in the FLN5+31 unfolded state ensemble after reweighting with the PRE NMR data. Cα, P, N3, and C4’ atoms and a distance cut-off of 12.5 Å were used to define NC-ribosome contacts. rRNA helices and ribosomal proteins surrounding the exit tunnel are annotated. **f,** Circos plot showing the interactions between the ribosome-anchored sub-ensemble of the FLN5+31 unfolded state (after reweighting with the PRE NMR data) and the ribosomal proteins and rRNA. The greyscale lines connecting the FLN5 NC and the ribosomal proteins and rRNA represent the extent of intermolecular interactions (% of total frames). The anchored sub-ensemble was defined using a -5 kJ mol^-1^ interaction energy cut-off between a previously identified C-terminal binding segment of FLN5 (residues 728-747) and the ribosome (Extended Data Figure 5c)^35^.

Consistent with the two distinct NC exit pathways identified within the ribosomal exit tunnel^34^, upon emergence the FLN5 ensemble predominantly interacts with ribosomal proteins uL23 and uL29 and 23S rRNA helix H7, whereas the A3A3 ensemble shows a preferred orientation toward ribosomal protein uL24 and 23S rRNA helices H24, H50 and H59 (Figures **2d**, **3e** and Extended Data Figure **5d**)^35,52^. In addition to the strong C-terminal binding segment spanning strands F, F’ and G, the PRE reweighted ribosome-anchored FLN5 sub-ensemble also exhibits transient ribosome interactions with strands A and A’ (Figure **3f**). By contrast, the PRE reweighted A3A3 ribosome-anchored sub-ensemble engages in transient ribosome interactions not only via strands A and A’, but also with strands D and E (Extended Data Figure **8c**).

Overall, strands C and C’ form non-native contacts within the FLN5 ribosome-anchored sub-ensemble, strands A and A’ engage in non-native contacts and transiently interact with the ribosome, and strands F, F’ and G form stable interactions with the ribosome surface. Native-like contacts are sparsely populated and only very transiently explored by strands A, B, C and E, suggesting that the combined effect of non-native interactions and ribosome binding contributes to delayed folding within the most populated wild-type FLN5 unfolded state sub-ensemble, although their relative contributions remain to be fully deconvoluted.

## Ribosome unbinding promotes an expanded unfolded state resembling the folding-incompetent A3A3 variant

Ribosome unbinding of the C-terminal segment of FLN5 (residues 728-747) gives rise to an expanded unfolded state, forming the second most populated FLN5 sub-ensemble on the ribosome (Figure **4a**).

**Figure 4.**
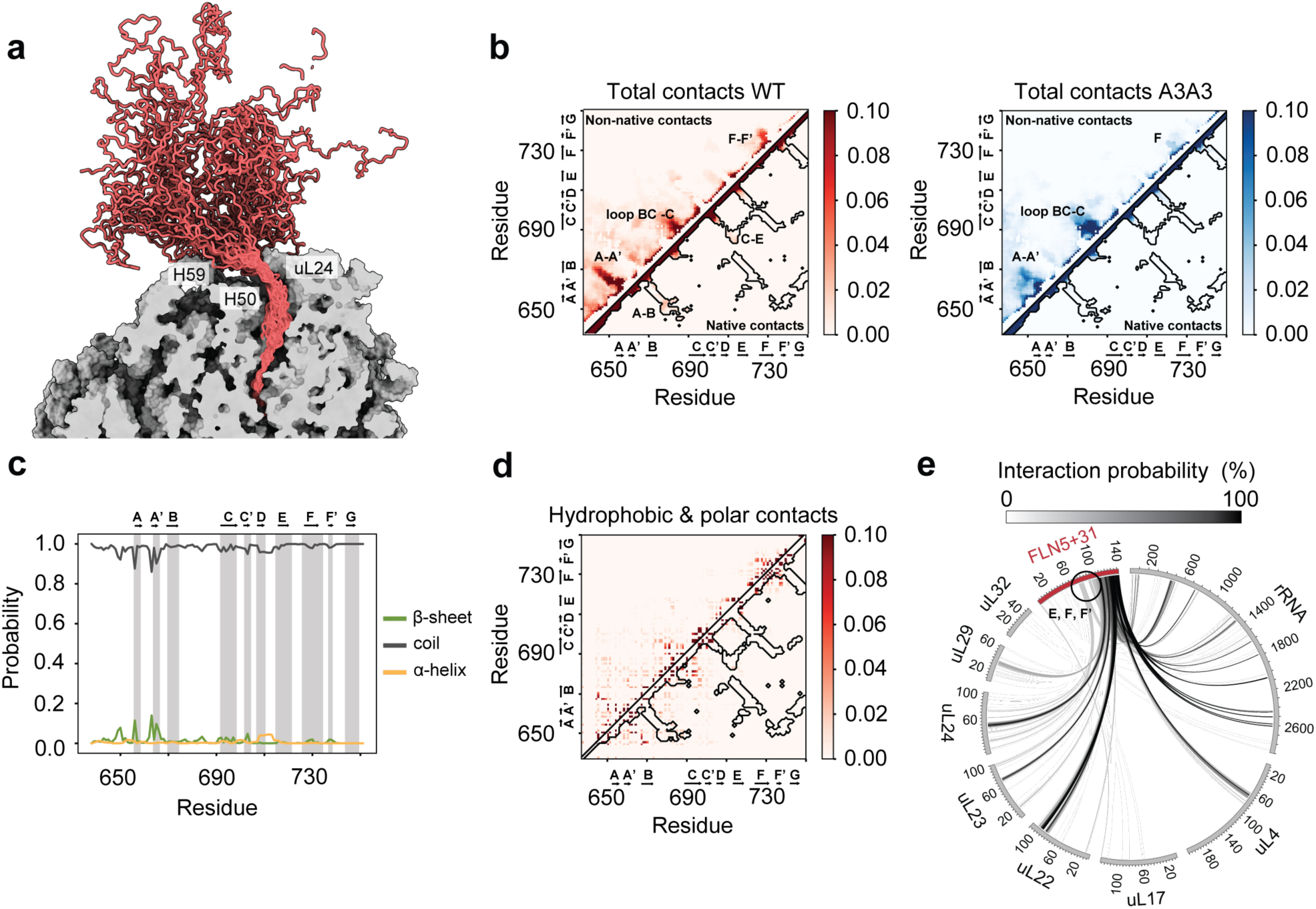
Structural preferences of the expanded sub-ensemble of the FLN5 unfolded state on the ribosome. **a**, A representative sub-ensemble (top 50 cluster centroids after reweighting with the PRE NMR data) of the expanded unfolded state of wild-type FLN5 on the ribosome. The ribosome cross-section is shown in grey and is sliced to enable visualisation of the FLN6 linker within the tunnel. **b,** Average contact maps of the expanded sub-ensemble of the unfolded state on the ribosome (zoomed in to a probability of 0.1) indicating total contacts in the wild-type FLN5+31 (left) and A3A3+31 (right), as well as contacts between hydrophobic and polar residues in the wild-type FLN5+31 **(d)**. Contacts were defined as Cα-Cα distances of less than 10Å and recapitulated when using closest heavy atom definitions (cut-off 6 Å; Extended Data Figure 9b). Non-native contacts are mapped above the diagonal; native contacts are mapped below the diagonal in **(b)**. The black contours highlight the native contact map of the folded FLN5 domain. **c,** Average secondary structure propensities in the expanded sub-ensemble of the wild-type FLN5 unfolded state on the ribosome along the protein sequence determined using the DSSP algorithm. The vertical shaded areas highlight the regions of β-strands (annotated as strands A-G) in natively folded FLN5. **e,** Circos plot showing the interactions between the expanded sub-ensemble of the FLN5+31 unfolded state (after reweighting with the PRE NMR data) and the ribosomal proteins and rRNA. The greyscale lines connecting the FLN5 NC and the ribosomal proteins and rRNA represent the extent of intermolecular interactions (% of total frames).

The expanded sub-ensemble comprises 13% of the wild-type FLN5 unfolded state on the ribosome and exhibits an average Rg of 46.2 ± 7.0 Å in FLN5 (Figure **2b**). It explores local non-native contacts similar to those observed in the ribosome-anchored sub-ensemble, including interactions between strands A and A’ at the N-terminus and between the loop connecting strands B and C, and strand C. The A-A’ contacts, while still of low occurrence (∼6-10%), are more prominent than in the ribosome-anchored sub-ensemble. In addition, this expanded sub-ensemble explores non-native contacts between strands F and F’, and transient native-like contacts (0.4 ± 0.1%) between strands C and E (Figure **4b**). These contacts, like those observed in the ribosome-anchored sub-ensemble, form predominantly between polar and hydrophobic residues (Figures **4d** and Extended Data Figure **9a**). The β-sheet propensity (1.3 ± 0.2%) is most pronounced for strands A and A’, as well as for a region in the N-terminal unstructured loop (Figure **4c**).

The A3A3 expanded sub-ensemble is seen to closely resemble the wild-type, sharing key local non-native contacts involving strands A and A’, and the loop connecting strands B and C, and strand C. However, it lacks the non-native contacts between strands F and F’ and the native contacts between strands C and E observed in the wild-type. Instead, it forms additional non-native contacts involving strand F alone (Extended Data Figure **8a**). The average β-sheet propensity is somewhat reduced (1.1 ± 0.2%) compared to wild-type and is best defined in the loop region connecting strands B and C, and in strand C (Extended Data Figure **8b**).

The expanded FLN5 sub-ensemble, by definition, lacks the stable ribosome interactions mediated by the C-terminal binding segment of FLN5 (residues 728-747). Nonetheless, in the PRE reweighted wild-type sub-ensemble, transient ribosome interactions are observed via strands E, F, F’ and G (Figure **4e**). The A3A3 sub-ensemble shows a similar pattern of transient ribosome interactions, but these do not involve strand E (Extended Data Figure **8c**).

Even after ribosome unbinding and the consequent loss of stable C-terminal interactions, the expanded FLN5 sub-ensemble does not explore native-like contacts to a significant extent. Instead, it closely resembles the A3A3 expanded sub-ensemble and likely remains unfolded through a concerted interplay of non-native contacts between strands A and A’, between the loop connecting strands B and C, and strand C, and between strands F and F’, alongside transient ribosome interactions involving strands E, F, F’ and G, while only strands C and E engage in transient native-like contacts.

## The least-populated compact sub-ensemble forms native-like contacts and seeds the co-translational folding nucleus

A compact sub-ensemble (Figure **5a**) is also observed and is the least populated in both the wild-type and A3A3 unfolded ensembles (Figure **2b**) yet is more than two-fold more populated in the folding-competent wild-type (9.7%) than in the folding-incompetent A3A3 variant (4.6%). The wild-type compact sub-ensemble exhibits an average Rg of 29.7 ± 3.7 Å (Figure **2b**), and shows a notable increase in native contacts (1.7 ± 0.1%) relative to the ribosome-anchored (0.6 ± 0.1%) and expanded (0.4 ± 0.1%) sub-ensembles, specifically between strands A and E, B and E, B and D, C and E, and C and F (Figures **5b**, **5e**).

**Figure 5.**
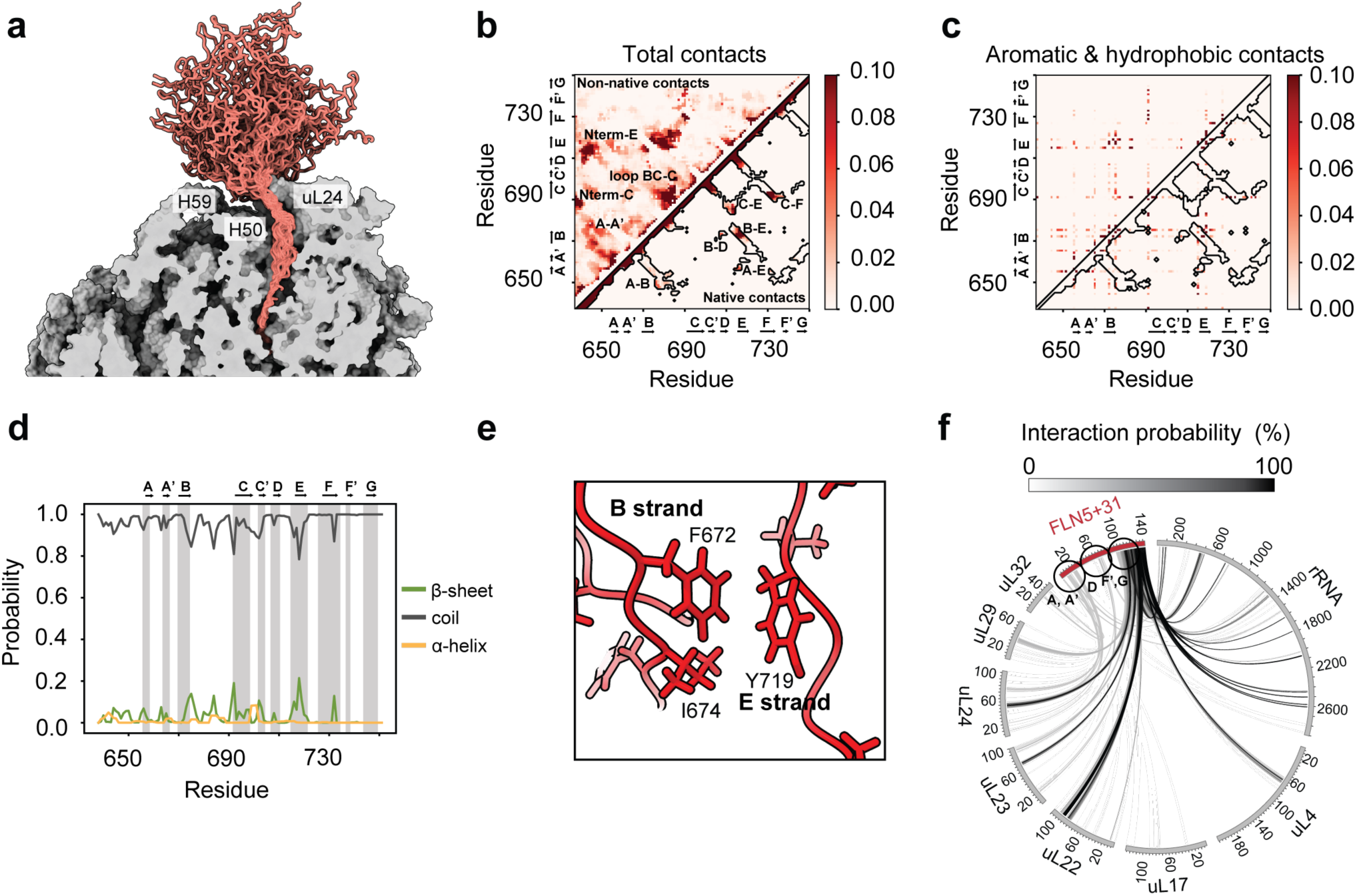
Structural preferences of the compact sub-ensemble of the FLN5 unfolded state on the ribosome. **a**, A representative sub-ensemble (top 50 cluster centroids after reweighting with the PRE NMR data) of the compact unfolded state of wild-type FLN5 on the ribosome. The ribosome cross-section is shown in grey and is sliced to enable visualisation of the FLN6 linker within the tunnel. Average contact maps of the compact sub-ensemble of the wild-type FLN5 unfolded state on the ribosome (zoomed in to a probability of 0.1) indicating total contacts **(b)** and contacts between hydrophobic and polar residues only **(c)**. Contacts were defined as Cα-Cα distances of less than 10Å and recapitulated using closest heavy atom contact definitions (cut-off 6 Å; Extended Data Figure 9b). Non-native contacts are mapped above the diagonal; native contacts are mapped below the diagonal in **(b)**. The black contours highlight the native contact map of the folded FLN5 domain. **d,** Average secondary structure propensities in the compact sub-ensemble of the wild-type FLN5 unfolded state on the ribosome along the protein sequence determined using the DSSP algorithm. The vertical shaded areas highlight the regions of β-strands (annotated as strands A-G) in natively folded FLN5. **e,** Representative intramolecular interactions within the compact sub-ensemble depicting a transiently formed near-native β-sheet between strands B and E stabilised by hydrophobic interactions between F672/I674 (strand B) and Y719 (strand E). **f,** Circos plot showing the interactions between the compact sub-ensemble of the FLN5+31 unfolded state (after reweighting with the PRE NMR data) and the ribosomal proteins and rRNA. The greyscale lines connecting the FLN5 NC and the ribosomal proteins and rRNA represent the extent of intermolecular interactions (% of total frames).

These native contacts predominantly form between hydrophobic and aromatic residues, consistent with the formation of an early folding nucleus (Figures **5c**, **5e** and Extended Data Figure **9a**). In addition, the sub-ensemble retains non-native contacts between strands A and A’ and between the loop connecting strands B and C, and strand C (Figure **5b**), involving interactions between hydrophobic and polar residues (Extended Data Figure **9a**), as previously observed in the ribosome-anchored and expanded sub-ensembles. It also explores additional non-native contacts between the N-terminal unstructured region and strands C and E (Figure **5b**), primarily involving polar and charged residues (Extended Data Figure **9a**). The compact sub-ensemble also exhibits approximately two-fold higher β-sheet propensity (2.8 ± 0.4%) compared to both the ribosome-anchored (1.5 ± 0.2%) and expanded (1.3 ± 0.2%) sub-ensembles, aligning with the positions of strands A, A’, B, C, C’, D, E and F, while the loop connecting strands B and E adopts a transient non-native β-sheet secondary structure (Figure **5c**).

The compact sub-ensemble of the folding-incompetent A3A3 variant explores fewer native-like contacts (1.2 ± 0.1%) compared to the wild-type, but retains those between strands A and E, B and D, and B and E, albeit to a lesser extent (Extended Data Figure **8a**). The non-native contacts between strands A and A’ are preserved, while additional non-native interactions emerge, including contacts between strand A’ and the loop connecting strands B and C, between strands A’ and F, F and F’, and between the N-terminal unstructured region, including the His-tag, and strand G (Extended Data Figure **8a**). The β-sheet propensity is somewhat reduced in A3A3 (2.0 ± 0.3%) and more pronounced for strands A’, B, C’, D and E, while strands F and F’ adopt a transient non-native β-sheet and an α-helix (Extended Data Figure **8b**).

The wild-type compact sub-ensemble engages in transient ribosome interactions via strands A, A’, D, F’ and G. Strand F no longer interacts with the ribosome, in contrast to both the ribosome-anchored and expanded sub-ensembles (Figure **5f**), consistent with its engagement in forming the coTF nucleus. In comparison, the corresponding A3A3 sub-ensemble engages the ribosome through strands E and F, as well as the N-terminal unstructured region, does not engage strand D, and maintains transient interactions via strands A’, F’, and G (Extended Data Figure **8c**).

Together, these results identify a compact sub-ensemble within the wild-type FLN5 unfolded state on the ribosome that is structurally distinct from both the ribosome-anchored and expanded sub-ensembles, as well as the compact sub-ensemble observed for the folding-incompetent A3A3 variant. It exhibits increased β-sheet propensity and forms early native-like contacts between strands A and E, B and E, B and D, C and E, and C and F. The N-terminal region, including strands A and A’, engages in non-native contacts, while strands A, A’, D, F’ and G interact transiently with the ribosome, suggesting that the wild-type compact sub-ensemble captures features characteristic of early folding events. Although folding nuclei are typically characterised by φ-value analysis during refolding^55–57^, PRE NMR performed at equilibrium can, in principle, detect low-population, high-energy states that exchange rapidly (k_ex_ > 1000 s^−1^) with the unfolded state^58–60^. In contrast, more structured intermediates exchange slowly (k_ex_ < 1 s^−1^)^34^ and remain PRE-invisible. We therefore suggest that the compact sub-ensemble provides structural insight into a transient, sparsely populated state that is likely reminiscent of the coTF nucleus.

## Discussion

The role of the unfolded state is contentious in the protein folding field ^31,61^. While it is broadly recognised to influence folding pathways and energetics^39^, its precise nature remains elusive, with most studies focused on denatured states^14,62–64^ or, at best, unfolded states under folding conditions^12,65^, with considerable variation in their compactness and residual structure^23,24,31^.

Here, we present the first atomistic model of the FLN5 unfolded state on the ribosome under folding conditions (Figure **2**), providing structural and mechanistic insight into the initiation of *de novo* protein folding during biosynthesis on ribosomes (Figure **6**). At this point of elongation, the entire domain has emerged from the ribosomal exit tunnel and is available to fold, yet the NC populates a predominant unfolded state (44 ± 8%), two partially structured intermediates I1 (25 ± 9%) and I2 (21 ± 8%), and a sparse native state (10 ± 4%)^34^, in accord with the altered energetics of NC states relative to those in the corresponding isolated polypeptide^39,40^. The FLN5 unfolded state on the ribosome populates three structurally distinct sub-ensembles defined by their degree of compaction and engagement with the ribosome (Figure **2, 6**)^35,39^. Although our experiments were performed at equilibrium and cannot resolve the kinetics of exchange between sub-ensembles, the progression from ribosome-anchored via expanded to compact sub-ensembles provides the basis for a tentative mechanistic inference, whereby loss of ribosome interactions leads to NC expansion and increased solvation, while its subsequent compaction is accompanied by the release of water molecules, entropically favourable for folding initiation (Extended Data Figure **5f-g**).

**Figure 6.**
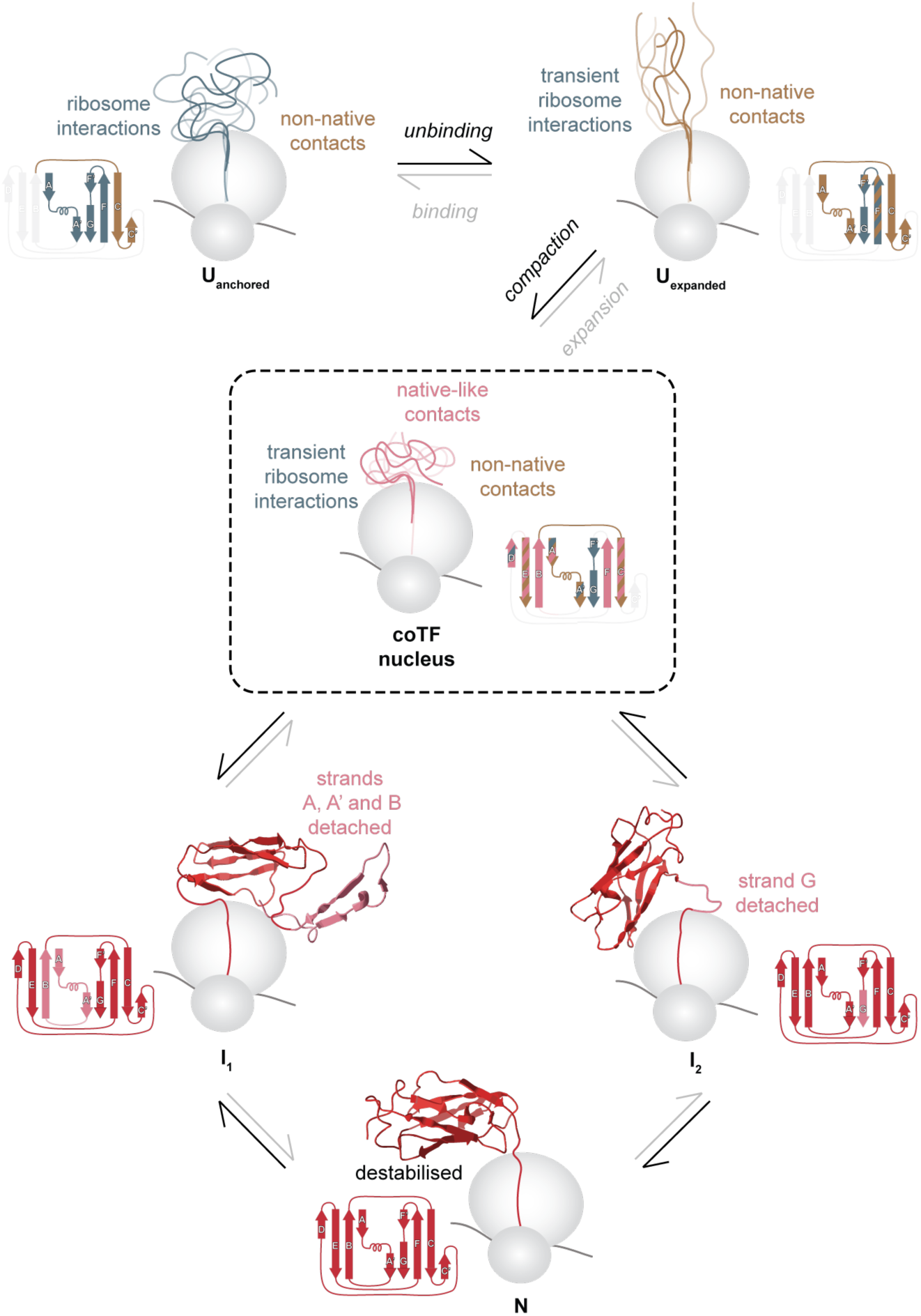
Model for the mechanism of *de novo* protein folding initiation on the ribosome at equilibrium. Progressive structure acquisition is illustrated using the topology representation of the native FLN5 domain with its secondary structural elements (PDB 6G4A^51^). Light grey denotes non-interacting disordered regions; blue grey denotes ribosome interactions; gold denotes non-native contacts; pink denotes native-like contacts and structure; red denotes the native structure. The nascent chain is coloured by the predominant interaction type within each sub-ensemble. U_anchored_: ribosome-anchored unfolded sub-ensemble; U_expanded_: expanded unfolded sub-ensemble; coTF nucleus: co-translational folding nucleus seeded by the compact unfolded sub-ensemble; I_1_ and I_2_: co-translational folding intermediates; N: native state.

The dominant sub-ensemble remains ribosome-anchored (Figure **3**), and is defined by local non-native contacts between strands A and A’, and between the loop connecting strands B and C, and strands C and C’, as well as stable ribosome interactions involving strands F, F’ and G, and transient ribosome contacts via strands A and A’. Only limited native-like contacts between strands B and E are transiently sampled, suggesting that not only ribosome-binding^35,66,67^, but also local non-native contacts can act to delay folding even after the full sequence of the FLN5 domain has emerged outside the ribosome^42^.

Upon ribosome unbinding, the unfolded NC populates an expanded sub-ensemble that is conformationally extended and predominantly non-native (Figure **4**). Although stable ribosome interactions are lost in this state, transient contacts persist via strands E, F, F’ and G. Non-native contacts between strands A and A’, and between the loop connecting strands B and C, and strand C, are maintained, while contacts involving strand C’ are no longer observed and new non-native contacts emerge between strands F and F’. Native-like contacts are rare and limited to transient interactions between strands C and E. Thus, non-native contacts in both the ribosome-anchored and expanded states, together with ribosome interactions, likely act to delay *de novo* folding by reshaping the coTF landscape, consistent with studies showing that non-native contacts can actively modulate folding pathways^26,28,30^. Correlation analysis suggests that in the ribosome-anchored sub-ensemble, a subset of non-native contacts show positive correlations with both N- and C-terminal ribosome interactions, while in the expanded sub-ensemble, positive correlations persist between non-native contacts and transient C-terminal ribosome interactions, consistent with a coordinated rather than independent modulation of the early folding landscape (Extended Data Figure **9c**)^26^.

Ultimately, the described entropic destabilisation of the expanded unfolded state^39^ leads to the formation of a structurally distinct compact sub-ensemble (Figure **5**). While non-native contacts persist in the N-terminal region, particularly between strands A and A’, and transient ribosome interactions involve strands A, A’, D, F’ and G, this sub-ensemble exhibits increased β-sheet propensity and is enriched in early native-like contacts between strands A and E, B and E, B and D, C and E, and C and F, consistent with the observation that native and non-native contacts act in concert during early folding events^26,28–30,68^. The coexistence of non-native and native-like contacts within the compact sub-ensemble suggests that non-native contacts do not simply delay folding but may actively facilitate the formation of the coTF nucleus by lowering the folding barrier for early folding events^69^. Thus, while non-native contacts likely delay folding in the ribosome-anchored and expanded sub-ensembles, their persistence into the compact sub-ensemble suggests a more nuanced role in shaping the landscape that ultimately enables the first native-like contacts to form. Furthermore, in the compact sub-ensemble, a subset of native-like and non-native contacts show positive correlations while ribosome interactions are partially negatively correlated with native-like contact formation, suggesting that release from the ribosome is a necessary step for folding initiation, rather than the resolution of all non-native contacts (Extended Data Figure **9c**)^26^.

Together, these observations indicate that the compact sub-ensemble populates a state reminiscent of the coTF nucleus (Figure **6**). The refolding of Ig-like domains has been extensively studied in isolation, where a nucleation-condensation mechanism has been proposed, driven by hydrophobic collapse and centred around native contacts between four conserved residues in strands B, C, E and F^55,70–75^. Notably, the folding nucleus in isolation is more structured than the state observed on the ribosome and does not involve strands A or D during early folding^72^, suggesting that the initiation of *de novo* protein folding during biosynthesis may involve a distinct nucleation mechanism from that observed during refolding *in vitro*, potentially one that accommodates the formation of downstream partially structured intermediates.

Indeed, FLN5 has been shown to adopt two partially structured intermediates, one along each of two parallel coTF pathways, referred to as I1 and I2; both featuring a structured core spanning strands C-F’: in I1, strands A, A’ and B are detached from the folded core, while in I2, only strand G remains dissociated (Figure **6** and Extended Data Figure **10a**)^34,48^. Although the compact sub-ensemble only transiently samples native-like contacts, they largely overlap with those present in both I1 and I2 (Extended Data Figure **10a-b**), consistent with the coTF nucleus seeding the formation of downstream partially structured intermediates (Figure **6**). While most non-native contacts are unique to the unfolded state sub-ensembles, the A-A’ non-native contacts observed in the expanded sub-ensemble partially overlap with those observed in I1 (Extended Data Figure **10b**). Notably, strands A and A’ also engage transiently with the ribosome surface in both the ribosome-anchored and compact sub-ensembles (Figures **3f**, **5f**), consistent with ribosome interactions observed at the unstructured N-terminal region of I1^48^. Together, this interplay between non-native contacts and ribosome interactions involving strands A and A’ across the unfolded sub-ensembles and into I1 suggests that both act in concert to maintain the N-terminal region unstructured along the coTF pathway. By contrast, strand G remains largely ribosome-anchored throughout all three unfolded sub-ensembles (Figures **3f**, **4e, 5f**), consistent with its dissociation from the folded core and ribosome interactions observed in I2^48^.

In conclusion, our findings establish the structural basis for a mechanistic model of *de novo* folding initiation during biosynthesis, revealing how a dynamic interplay between ribosome interactions, non-native contacts and early native-like contacts shapes the co-translational folding landscape from the earliest steps of structure formation. The initiation of *de novo* folding on the ribosome does not simply recapitulate the nucleation-condensation mechanism observed during refolding *in vitro*^55,70–75^, but proceeds via a distinct pathway in which the ribosomal environment actively modulates the energy landscape of the nascent chain. Together with our prior characterisation of the unique thermodynamics of coTF^39^ and the structures of downstream co-translational folding intermediates^48^, these findings complete an atomistic description of a co-translational folding pathway.

**Extended Data Figure 1.**
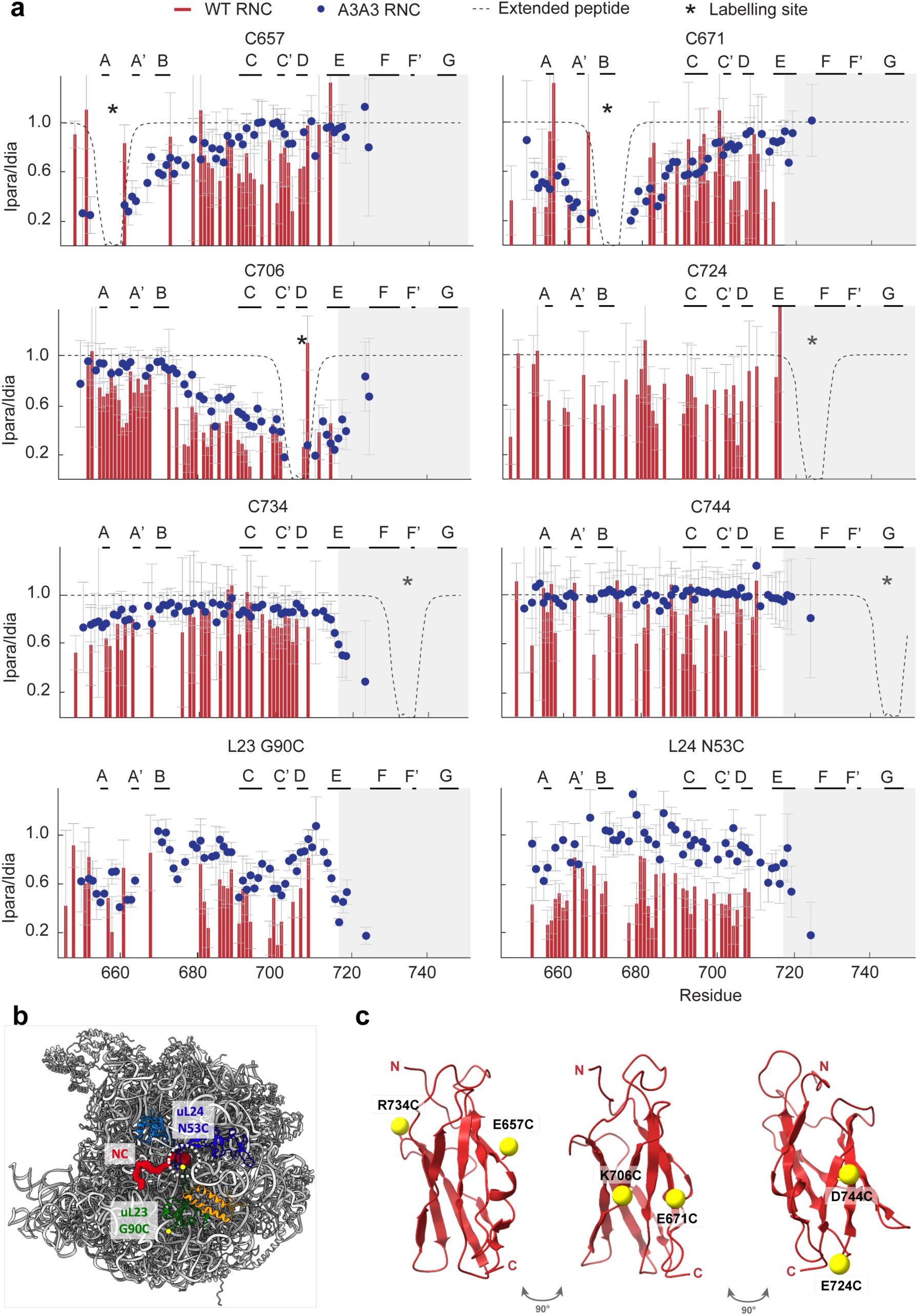
PRE analysis of unfolded wild-type FLN5 and A3A3 on and off the ribosome. **a**, PRE intensity ratio profiles for eight different labelling sites (in the nascent chain (NC) and on the ribosome; indicated with the black star) in the wild-type FLN5 (red) and A3A3 (blue) NCs on the ribosome. NMR data were recorded at 800MHz, 283K. Theoretical reference profiles expected for a fully extended polypeptide are also shown as dashed lines^39^. The secondary structure elements (β-strands) of native FLN5 are indicated at the top. The shaded region at the C-terminus represents the region of FLN5 that is broadening beyond detection through ribosome interactions (N730-K746, in the RNC). Errors were propagated from the standard deviation of the spectral noise. The second panel with grey bars under each dataset shows the difference between the RNC and isolated data. **b,** Annotated MTSL labelling sites (yellow circles) on the ribosome structure near the exit tunnel. **c,** The annotated crystal structure (PDB 1QFH (108)) is shown from three views, highlighting the PRE labelling sites used for RNC labelling.

**Extended Data Figure 2.**
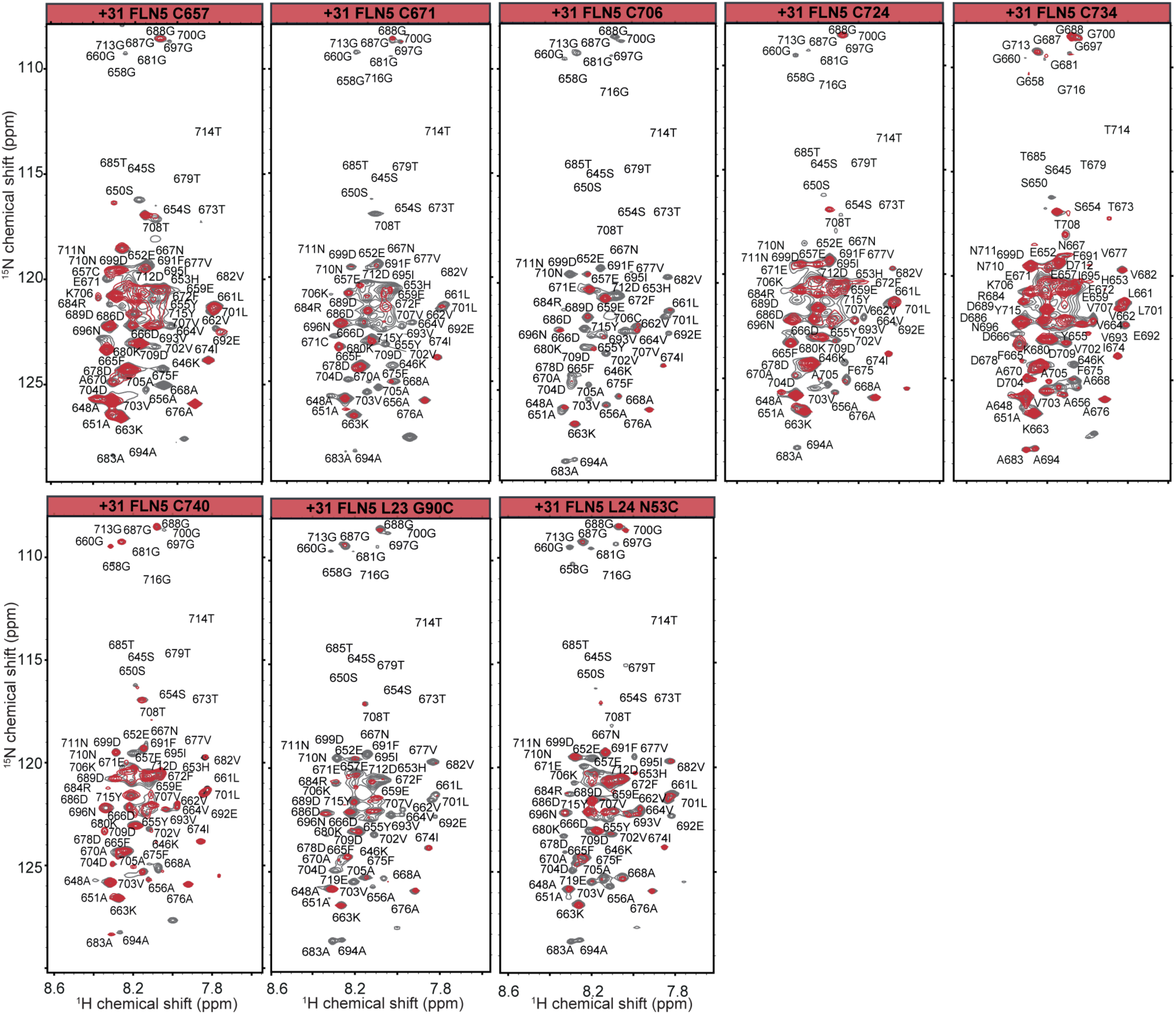
^1^H-^15^N SOFAST-HMQC PRE-NMR spectra of FLN5 RNC variants. The paramagnetic (red) and diamagnetic (grey) spectra are overlaid and residue assignments labelled. All spectra were recorded at 800MHz, 283K.

**Extended Data Figure 3.**
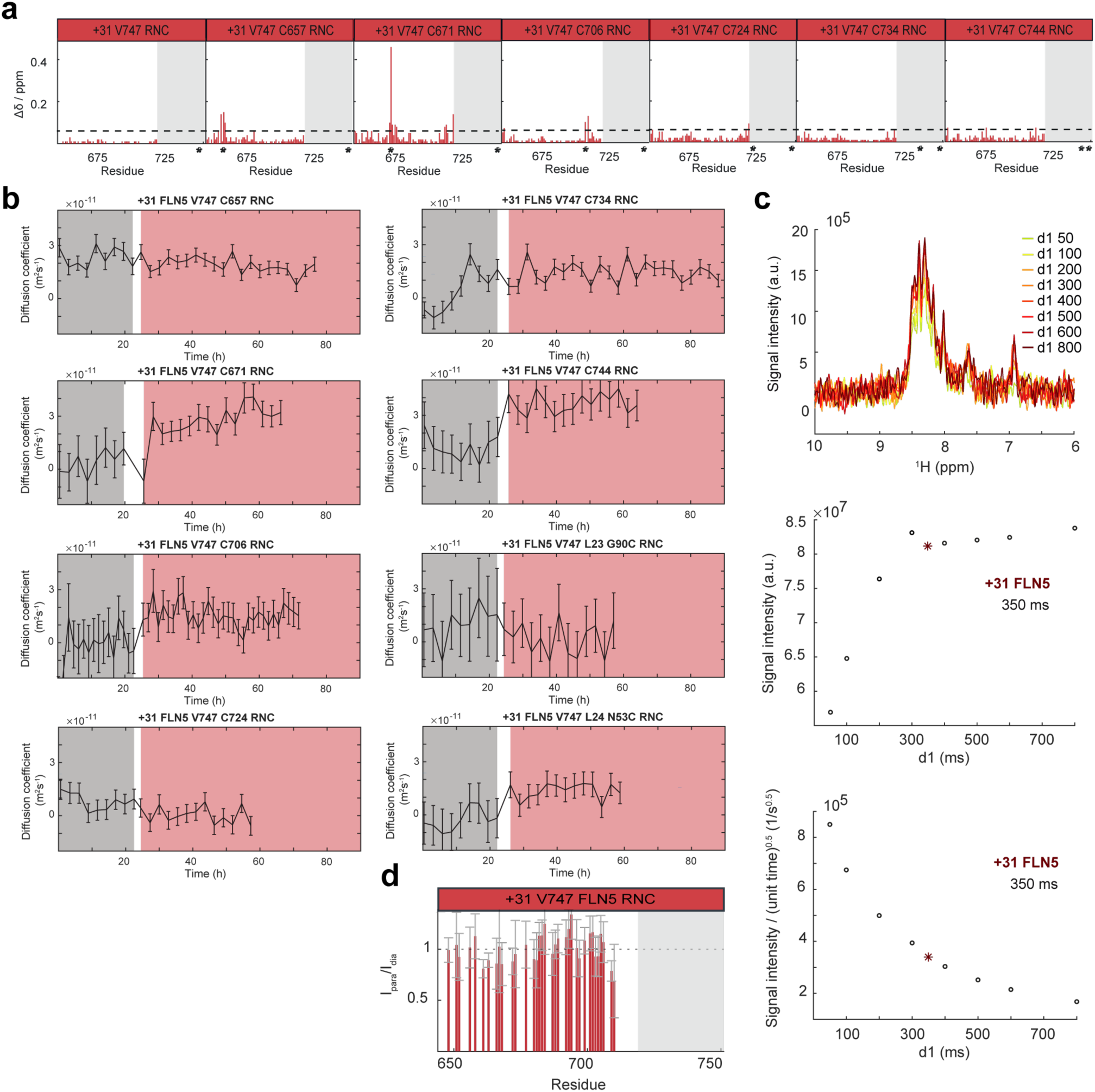
Quality control and optimisation of PRE-NMR experiments. **a**, Chemical shift perturbations (CSPs) along the protein sequence for all MTSL-labelled FLN5 RNC variants measured in the ^1^H-^15^N SOFAST-HMQC spectra of FLN5+31 RNC cysteine variants relative to the the FLN5+31 RNC. The labelling sites are indicated with a star (*). The dotted line indicates a threshold of 0.06ppm. **b,** Integrity of RNCs during PRE experiments was monitored with ^15^N-SORDID diffusion measurements. The calculated diffusion coefficient D is shown throughout NMR acquisition (centre), highlighting the paramagnetic (grey) and the diamagnetic acquisition timeframe (red). Error bars indicate propagated errors derived as standard deviation of the spectral noise. **c,** Optimisation of the recycle delay (d1) time chosen for PRE SOFAST-HMQC experiments to provide maximum sensitivity while also allowing the signal to relax completely before the subsequent scan is initiated. 1D ^1^H spectra at d_1_ values ranging from 50-800 ms (top, yellow to red gradient); total signal intensity dependence on the d_1_ value (middle); time-averaged signal (bottom). 350ms was chosen for PRE experiments. **d,** PRE intensity ratios of the FLN5+31 variant without any cysteines in the NC (C747V, Δcys). Error bars indicate propagated errors derived from the spectral noise.

**Extended Data Figure 4.**
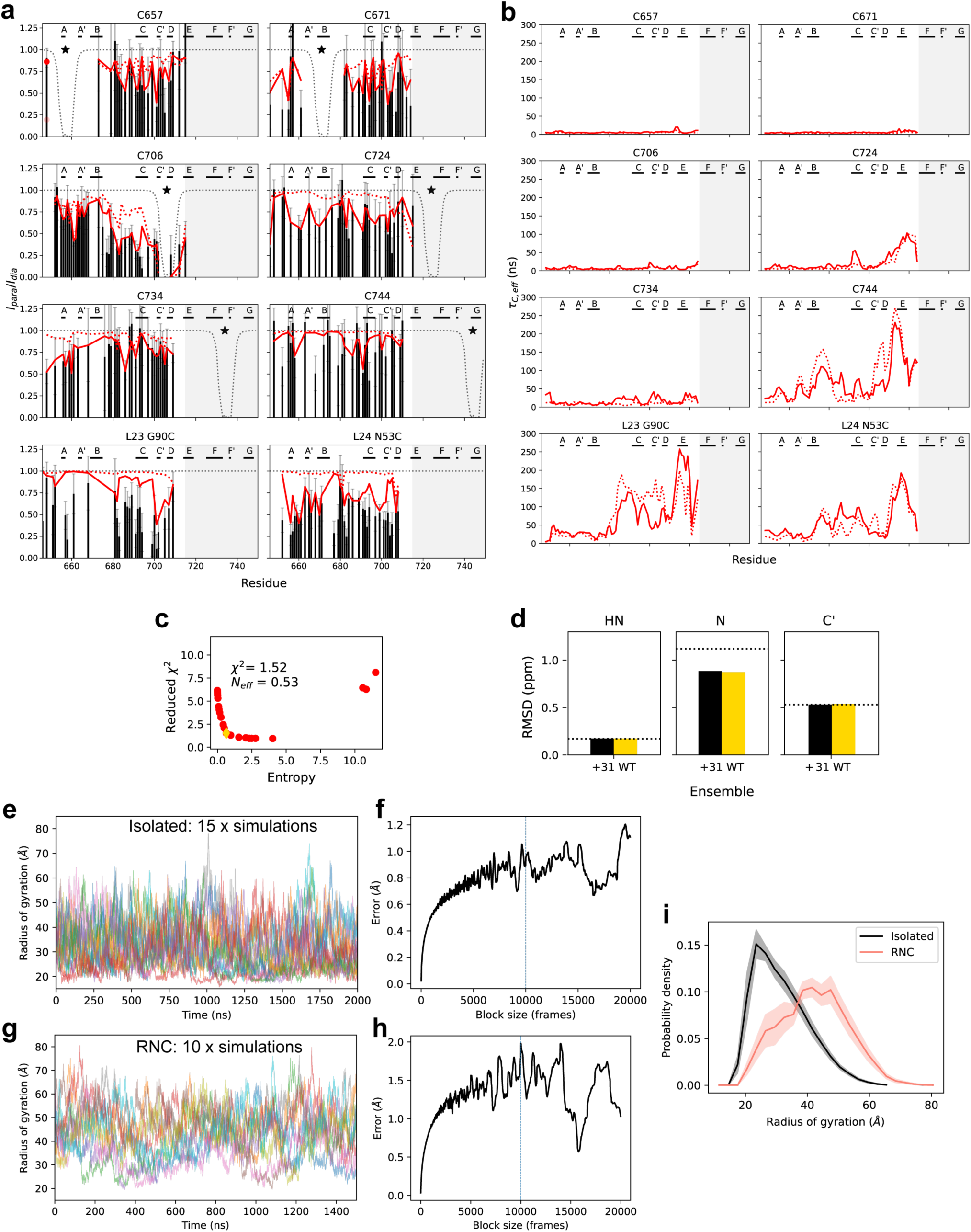
Bayesian reweighting with experimental PRE data and comparison to NMR backbone chemical shifts. **a**, Overlay of the experimental PRE data (black bars) and simulated PREs before (dotted line) and after (solid line) reweighting for FLN5+31. **b,** Analysis of the effective correlation times (τ_C,eff_) between the nitroxide labelling site and corresponding amide proton nuclei before (dotted line) and after reweighting (solid line) for FLN5+31. **c,** L-curve analysis of the reweighting calculations showing the agreement with the experimental data on the y-axis (χ^2^) and extent of fitting on the x-axis (entropy term, corresponding to the Kullback-Leibler divergence). The statistics at the elbow are indicated on the plots, including the effective fraction of frames left after reweighting (N_eff_). The simulated PREs in solid lines in **(a)** represent the elbow of the L-curves. **d,** Agreement with of the ensembles before (black) and after (yellow) reweighting with experimental chemical shifts. The dotted horizontal line indicates the average error of the forward model used to calculate the chemical shifts. **e-g,** Time-courses of the radius of gyration of the unfolded state in isolation (panel **e**) and as an RNC (panel **g**) observed in all-atom MD simulations. Each separate line/colour represents an independent trajectory. **f, h,** Block-averaging analysis showing converged error estimates of the Rg values presented in panels **e** and **g**, respectively. **i,** Rg probability distributions of isolated and RNC unfolded states with errors from blocking indicated with the shaded regions.

**Extended Data Figure 5.**
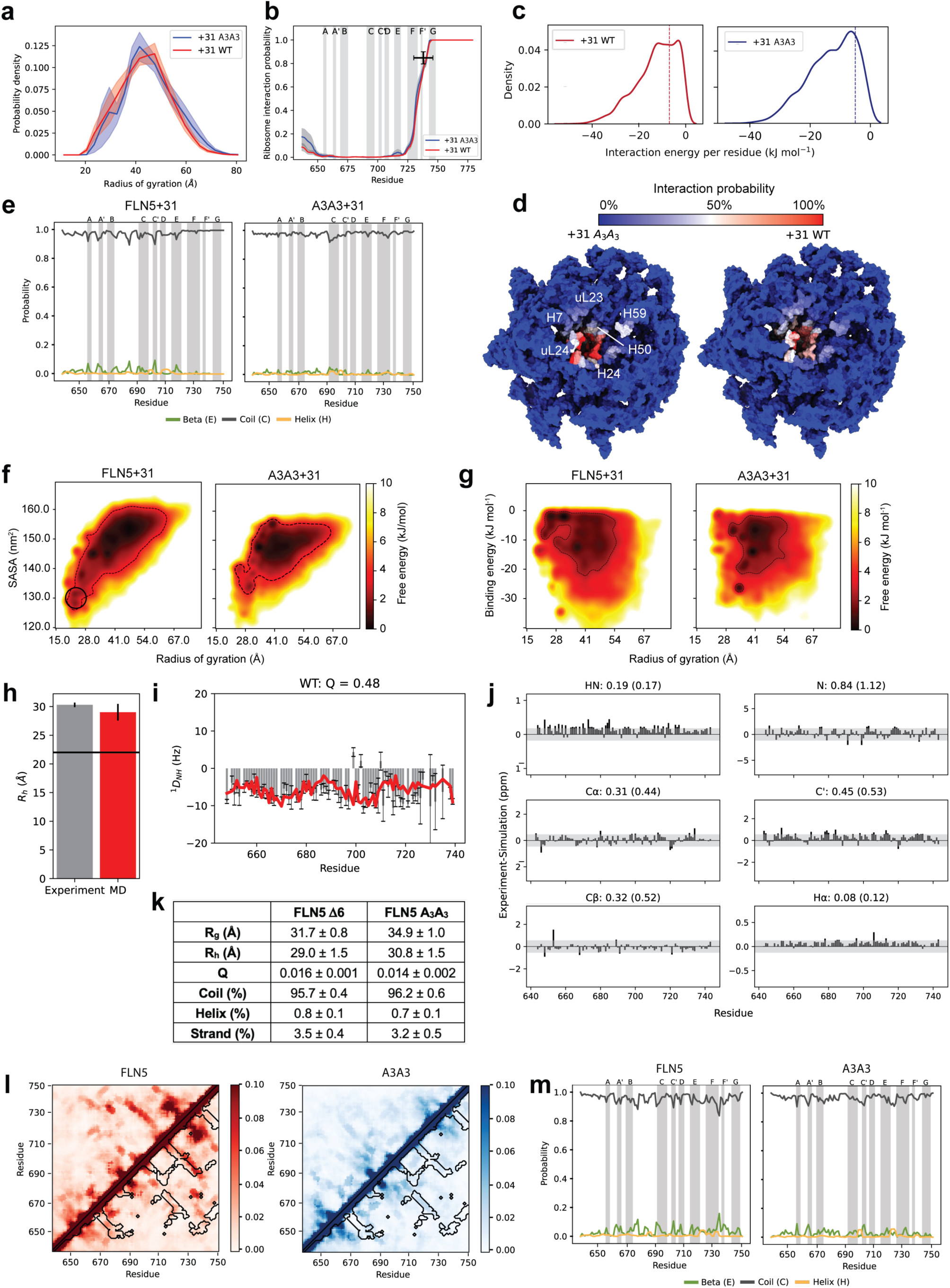
Comparison of the RNC and isolated polypeptide unfolded state ensembles of FLN5 and A3A3. **a**, Probability distributions of the all-atom radius of gyration (R_g_) for the FLN5 segment (residues 637-750). The error is estimated from block averaging. **b,** Average interaction probability between the nascent chain and ribosome surface plotted along the FLN5 sequence. The error is also estimated from block averaging and the secondary structure elements of FLN5 are highlighted. The black cross represents the experimentally measured interaction probability between the C-terminus and ribosome surface. The bar along the x-axis represents the binding region (residues 728-747) as defined previously. **c,** Probability distribution of the average interaction energy between the FLN5 C-terminus (residues N730-G750) and ribosome surface observed in the reweighted (with the PRE-NMR data) structural ensemble of FLN5+31 calculated using only NC and ribosome atoms with the C36m force field parameters. The vertical dashed line represents the chosen cut-off of -5 kJ mol^-1^ to distinguish the anchored and free ensembles. **d,** Interactions of the nascent chain with the ribosome mapped on the ribosome surface of the simulation model including the vicinity of the exit tunnel (see methods). The bottom surface represents the difference contact probability highlighting regions contacted more by A3A3+31 in blue and for FLN5+31 in red. Only Cα, P, N3, and C4’ atoms and a distance cut-off of 12.5 Å were used to define a NC-ribosome contact/interaction. The main structural elements including rRNA helices and ribosomal proteins around the exit tunnel are annotated. **e,** Average secondary structure propensities along the protein sequence determined using the DSSP algorithm. The vertical shaded areas highlight the regions of β-strands (annotated as strands A-G) in natively folded FLN5. **f**, Free energy landscapes of unfolded FLN5+31 WT (left) and A3A3 (right) as a function of radius of gyration (R_g_) and solvent-accessible surface area (SASA). The dashed black contour represents a ‘low-free energy region’ at 3.7 kJ/mol for the purpose of visualisation. The circle in the WT panel (left) highlights a low-energy, compact sub-ensemble that is not as favourable in A3A3 (right) and is referred to as the compact sub-ensemble. **g,** Free energy landscapes of unfolded FLN5 WT and A3A3 on the ribosome as a function of radius of gyration and ribosome interaction energy. The dashed black contour represents a ‘low-free energy region’ at 2.5 kJ/mol for the purpose of visualisation. **h,** Comparison of the experimental (30.3 ± 0.4Å, FLN5 Δ6 unfolded state) and simulated (29.0 ± 1.5Å, calculated for residues 646-744) radius of hydration (R_h_). The R_h_ of the native state (22.0 ± 0.1Å) is shown as a horizontal line. The error of the simulated value represents the error of the forward model used (see methods). **i,** Experimental (FLN5 Δ6 unfolded state measured in PEG-Octanol, 283K, grey bars) and simulated ^1^D_NH_ RDCs (red line) are shown and the Q-factor quantifying the agreement. **j,** Difference between experimental (FLN5 Δ6 unfolded state, 283K) and simulated NMR backbone chemical shifts. The transparent grey area indicates the error of the forward model used to back-calculate the chemical shifts (see methods). The average RMSD and error of the forward model are shown in the title of each respective plot. **k,** Ensemble-averaged properties of the unfolded state of FLN5 Δ6 and FLN5 A3A3. The errors were obtained from block averaging. **l,** The contact maps of the isolated unfolded FLN5 (red) and A3A3 (blue) ensembles are shown. The black contours represent the native contacts and the colour bars represent contact probability. **m,** Average secondary structure propensities along the protein sequence determined using the DSSP algorithm. The vertical shaded areas highlight the regions of β-strands (annotated as strands A-G) in natively folded FLN5.

**Extended Data Figure 6.**
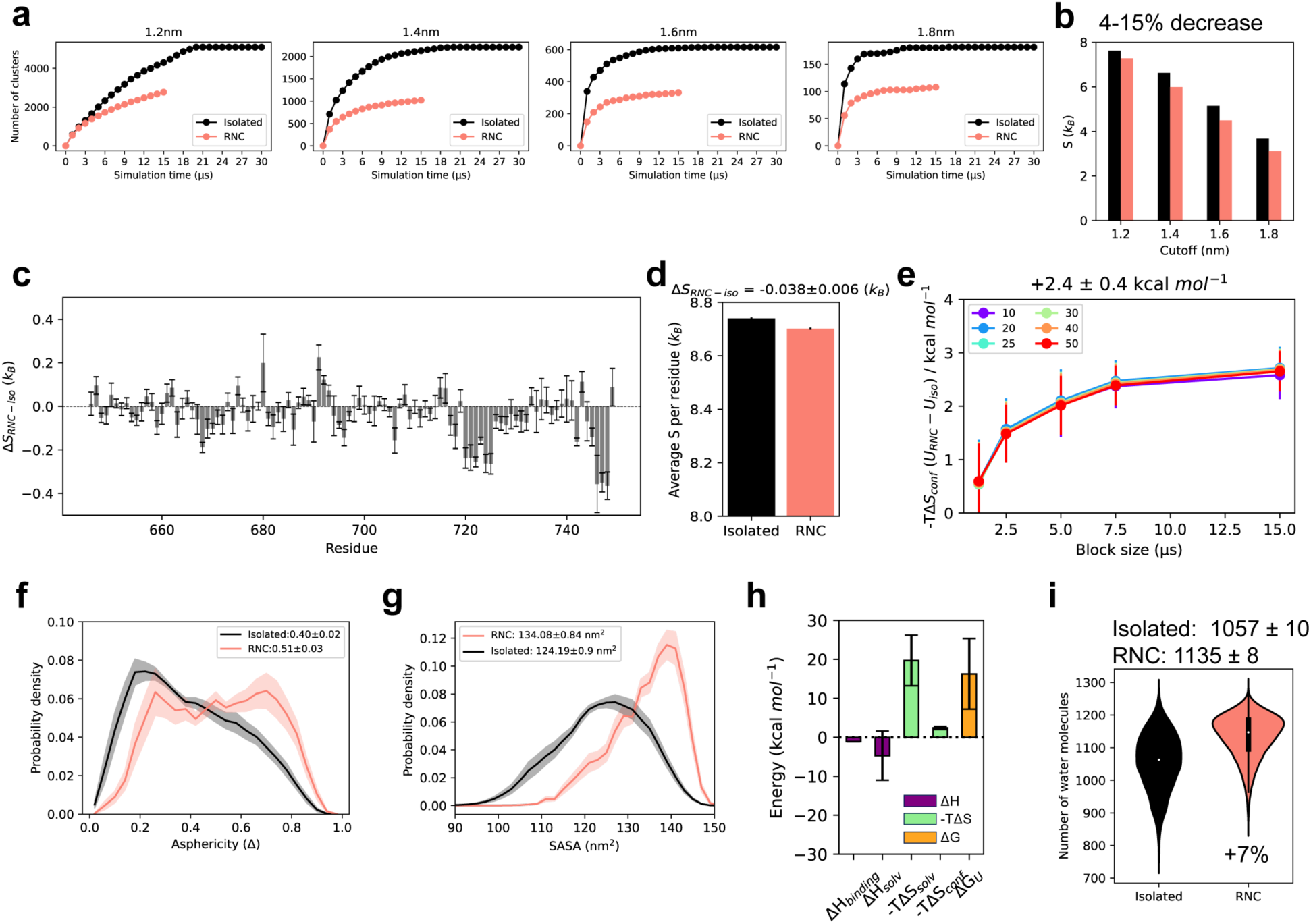
Analysis of the energetics of the wild-type unfolded state on and off the ribosome. **a**, Convergence of the number of clusters visited for several different cut-off values was assessed by plotting number of clusters as a function of simulation time^39^. This confirmed that sampling has been sufficient to reach a plateau in the number of clusters visited. This was analysed to ensure that differences between the RNC and isolated protein are not due to differences in sampling. **b,** The Gibbs entropy (− ∑*^n^ p_i_* × *ln*(*p_i_*), where n is the number of clusters/microstates and p the population of each microstate) was then estimated from the full ensembles after reweighting with the PRE data. **c,** Residue-specific total dihedral entropy^39^. The average and standard error obtained from block averaging are shown. **d,** The average S per residue residues for each ensemble is shown, using the average and standard error obtained from block averaging. **e,** The change in free energy (-TΔS = -TΔS_resi_N_resi_R) estimated from different block sizes of sampling indicates that the estimate, as expected, is dependent on sampling and appears to plateau at higher sampling times. The effect of the number of bins used to create the histograms (indicated in the legend) is minor. Thus, we consider this estimate a lower bound. **f,** The asphericity (Δ, see methods) probability distributions of the ensembles are shown (average and error displayed in figure legend). Errors were calculated using block averaging. **g,** Probability density function of the total solvent-accessible surface area (SASA) of FLN5 (residues 646-750) is shown for each ensemble with the average and standard error from block averaging displayed in the legend. Surfaces were calculated without the ribosome (−70S) and with the ribosome (+70S). The thermodynamic parameters of the solvation free energy difference between the unfolded state on and off the ribosome were calculated based on the apolar and polar changes in surface area and experimentally-parameterised functions of the heat capacity, C_p_, entropy, S, and enthalpy, H^39^. **h,** Summary bar plot of the enthalpic, entropic and total energetic changes on the ribosome (RNC-isolated). **i,** The number of water molecules contacting FLN5 (residues K646-G750) defined as a water center of mass to protein distance of less than 3.5Å. Water coordinates were saved every 1ns and all frames containing water from 10×3μs and 10×1.5μs, for the isolated and RNC ensemble respectively, were used for the analysis. The average and standard error obtained from block averaging is shown.

**Extended Data Figure 7.**
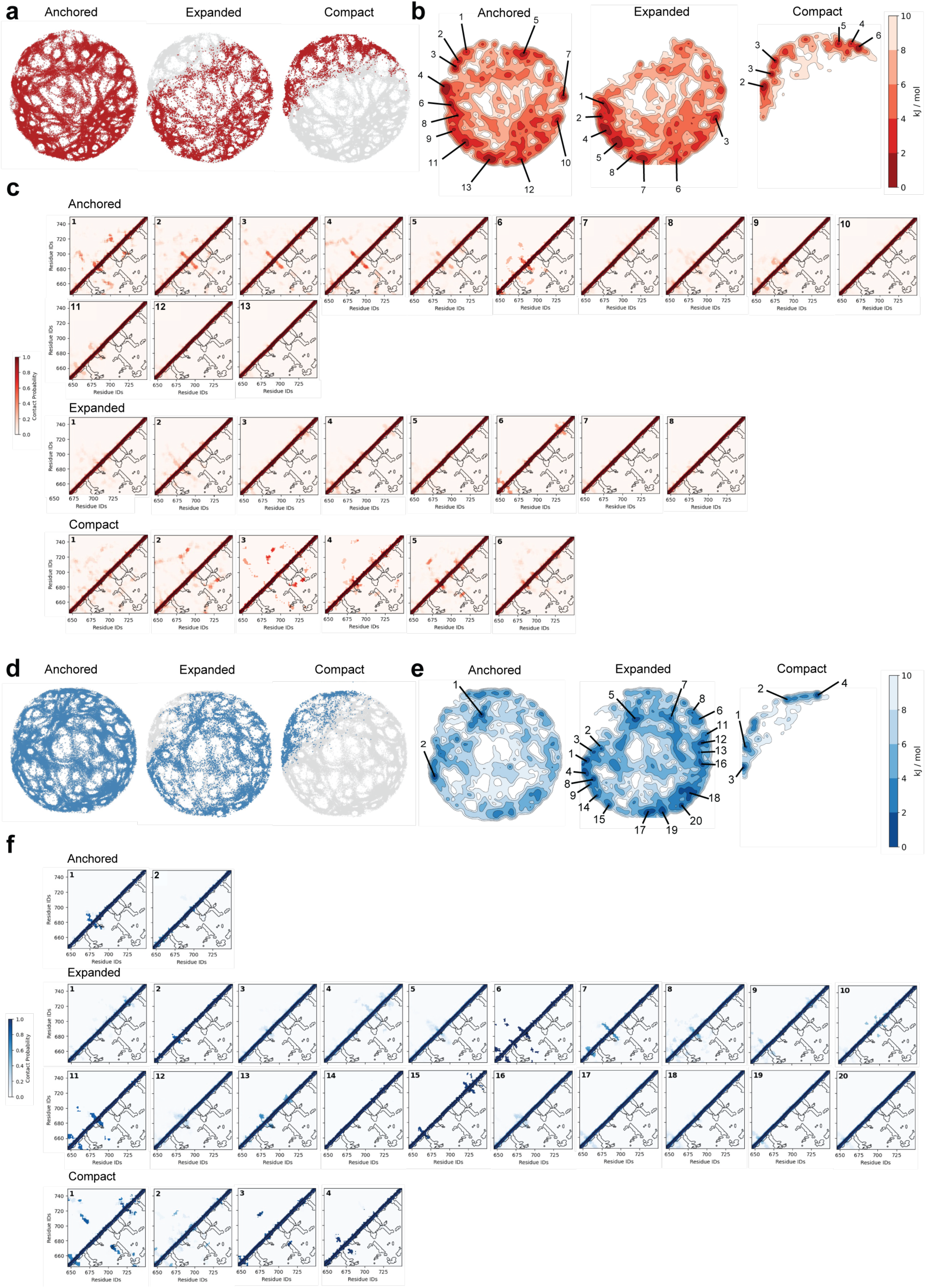
Identification of sub-ensembles on energy landscape visualisation method (ELViM) projections for WT FLN5 (red) and A3A3 ensembles (blue). **a**, Colouring of ELViM projection indices corresponding to frames from sub-ensembles (anchored, expanded, compact) from WT FLN5 ensemble. **b,** 2D free energy surface corresponding to density of ELViM projection of each sub-ensemble from the WT FLN5 ensemble. Frames from most populated minima (<2 kJ/mol) extracted for downstream analysis of contacts within sub-populations of each sub-ensemble, labelling mapped onto free energy surface to correspond to contact map labels. **c,** Average contact maps of most densely populated sub-populations of sub-ensembles identified from free energy surface of ELViM projection of WT FLN5. Black contours represent the native contact map of folded FLN5 and contacts were defined by 10.0 Å cutoff between Cα atoms. **d,** Colouring of ELViM projection indices corresponding to frames from sub-ensembles from A3A3 FLN5 ensemble. **e,** 2D free energy surface corresponding to density of ELViM projection of each sub-ensemble from the A3A3 FLN5 ensemble. Frames from most populated minima (<2 kJ/mol) extracted for downstream analysis of contacts within sub-populations of each sub-ensemble, labelling mapped onto free energy surface to correspond to contact map labels. **f,** Average contact maps of most densely populated sub-populations of sub-ensembles identified from free energy surface of ELViM projection of A3A3 FLN5. Black contours represent the native contact map of folded FLN5 and contacts were defined by 10.0 Å cutoff between Cα atoms.

**Extended Data Figure 8.**
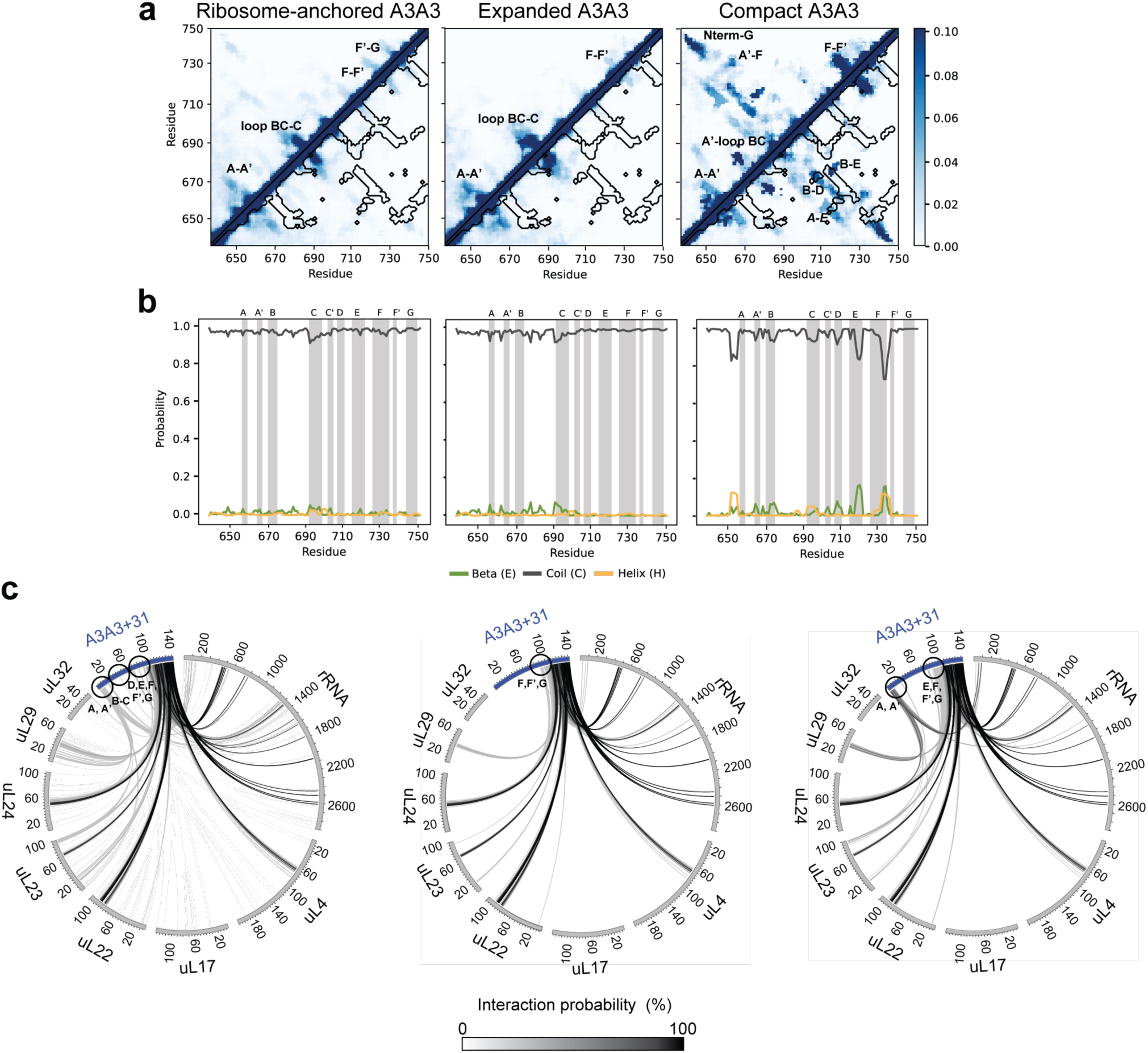
Analysis of ribosome-anchored (left), expanded (middle) and compact (right) sub-ensembles of the A3A3+31 unfolded state obtained from all-atom MD simulations after reweighting with PRE restraints. **a**, Average contact maps of the sub-ensembles (zoomed in to a probability of 0.1 for clarity). The black contours highlight the native contact map of folded FLN5. Cα atoms within 10.0 Å were used to define contacts. **b,** Average secondary structure propensities along the protein sequence determined using the DSSP algorithm. The vertical shaded areas highlight the regions of β-strands (annotated as strands A-G) in natively folded FLN5. **c,** Circos plots showing the interactions between the A3A3 NC and the ribosomal proteins and rRNA surrounding the exit tunnel as observed in the A3A3+31 unfolded state sub-ensembles. The greyscale lines connecting the A3A3 NC and the ribosomal proteins and rRNA represent the extent of intermolecular interactions (% of total frames).

**Extended Data Figure 9.**
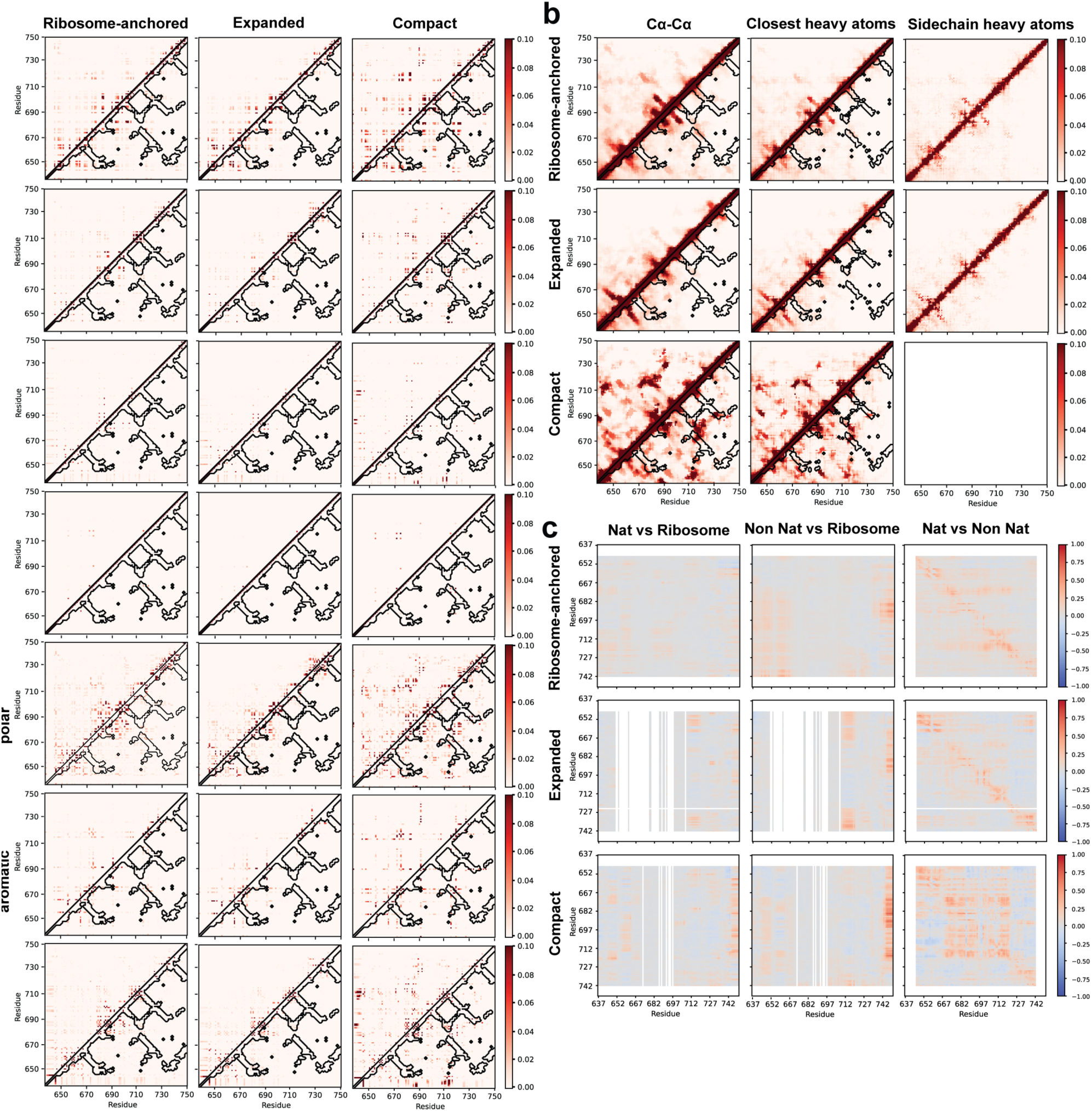
Analysis of the type, contact definitions and correlation between residues forming contacts in the ribosome-anchored (left), expanded (middle) and compact (right) sub-ensembles of the FLN5+31 unfolded state obtained from all-atom MD simulations after reweighting with PRE restraints. **a**, Average contact maps of the sub-ensembles (zoomed in to a probability of 0.1 for clarity). The black contours highlight the native contact map of folded FLN5. Cα atoms within 10.0 Å were used to define contacts. Residues were classified as follows: Hydrophobic (Ala, Val, Leu, Ile, Met, Pro), Polar (Ser, Thr, Cys, Asn, Gln, Gly), Positively charged (Arg, Lys, His), Negatively charged (Asp, Glu), Aromatic (Phe, Trp, Tyr). **b,** Average contact maps of the sub-ensembles (zoomed in to a probability of 0.1 for clarity). The black contours highlight the native contact map of folded FLN5. Cα-Cα contacts (left) defined with a distance cut-off of 10.0 Å, closet heavy (middle) and closest sidechain heavy contacts (right) defined with a distance cut-off of 6.0 Å. For Gly residues, the hydrogen atom is used to calculate closest sidechain heavy contacts. **c,** Plots capturing correlations between interchain native and non-native contacts, as well as NC-ribosome interactions for each NC residue in each sub-ensemble.

**Extended Data Figure 10.**
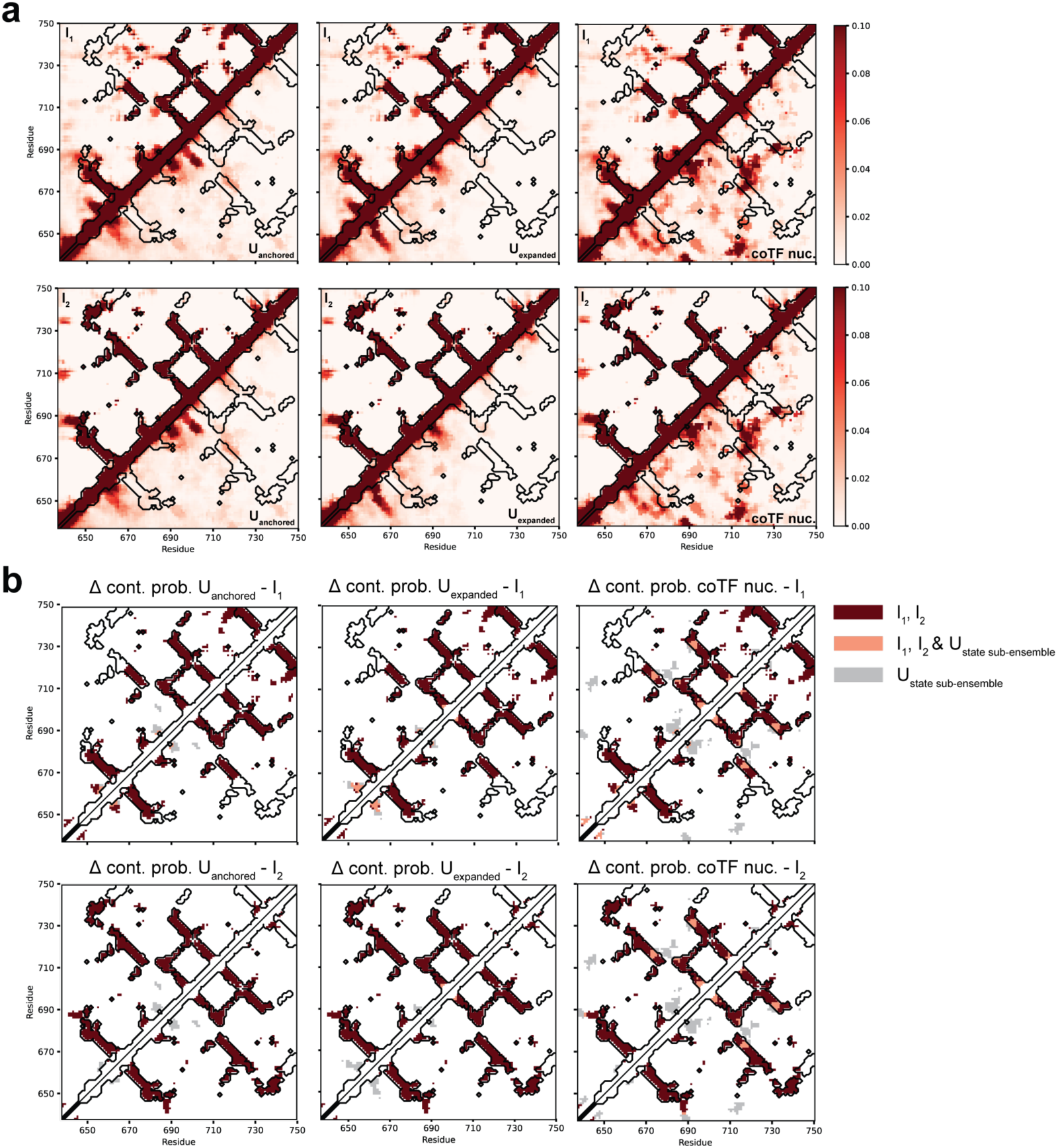
Comparison of the wild-type unfolded state sub-ensembles and the I1 and I2 folding intermediates. **a**, Average contact maps of the unfolded state sub-ensembles depicted below the diagonal: anchored (left), expanded (middle), compact (right), and intermediates depicted above the diagonal: I1 (top row) and I2 (bottom row). **b,** Difference contact probability maps of unfolded state sub-ensembles: anchored (left), expanded (middle), compact (right), and intermediates: I1 (top row) and I2 (bottom row). Dark red: contacts present in I1, I2; salmon: contacts present in unfolded state sub-ensembles and I1 and I2; grey: contacts present in the unfolded state sub-ensembles. In all panels, the contacts are zoomed in to a probability of 0.1 for clarity. The black contours highlight the native contact map of folded FLN5. Cα-Cα contacts are defined with a distance cut-off of 10.0 Å.

## Methods

### Protein expression and purification

DNA constructs of FLN5 were previously described^35,39,42^, and additional mutations were introduced via site-directed mutagenesis. FLN5 variants were expressed as His-tagged, isotopically labelled ribosome–nascent chain complexes (RNCs) in E. coli BL21 (DE3)-Gold cells, as previously described ^35,39^. All RNC constructs contained an arrest-enhanced variant of the SecM stalling sequence (FSTPVWIWWWPRIRGPP)^39^. Purification of isolated Δ6 FLN5 was carried out using affinity chromatography, followed by size exclusion chromatography in the presence of 6 M urea, prior to buffer exchange into Tico buffer (10 mM Hepes, 30 mM NH₄Cl, 12 mM MgCl₂, 1 mM EDTA).

For PRE-NMR experiments, the additional mutation C747V (referred to as FLN5 V747) was introduced, as previously described^39^. RNCs were uniformly labelled with ¹⁵N and purified as previously described^35,39^. For intermolecular PRE-NMR experiments, RNCs were expressed in E. coli BL21 strains harbouring CRISPR-engineered cysteine mutations in uL23 and uL24, as previously described^39,45^. All RNC samples were prepared in Tico buffer for experiments.

### NMR spectroscopy

NMR experiments of FLN5 were performed in Tico buffer at pH 7.5 and 283K. Chemical shifts were previously assigned and deposited^42^. All samples contained 10% (v/v) D2O and 0.001% (w/v) DSS as a reference. NMR data were recorded using a 700- (^1^H,^15^N, ^13^C NMR) or 800-MHz (^1^H,^15^N NMR) Bruker Avance III spectrometer, both equipped with a TCI cryoprobe, and Topspin 3.5pl2. Data were processed using NMRPipe (v11.1)^76^, Sparky (v3.12)^77^ and MATLAB (R2020b, The Mathworks Inc.).

For PRE-NMR experiments, we used a cysteine-less FLN5 C747V construct with six cysteine labelling sites introduced in the nascent chain and two on the ribosome surface. RNCs were reduced in Tico buffer supplemented with 2 mM TCEP, followed by buffer exchange into labelling buffer (50 mM HEPES, 12 mM MgCl₂, 20 mM NH₄Cl, 1 mM EDTA, pH 8.0), and labelled overnight at 277 K with a 10-fold molar excess of MTSL, as previously described^39^. Excess spin label was removed by buffer exchange into Tico. PREs were measured using 2D ¹H–¹⁵N SOFAST-HMQC experiments recorded at 800 MHz and 283 K, with acquisition times of 100 ms and 35 ms in the direct and indirect dimensions, respectively, and an inter-scan delay of 350 ms to allow full relaxation. Diamagnetic spectra were acquired following reduction with 2.5 mM sodium ascorbate. PREs were extracted by fitting cross-peaks to Lorentzian lineshapes in NMRPipe, applying 15 Hz and 5 Hz line-broadening in the direct and indirect dimensions, respectively, and converted to Γ₂ rates for Bayesian ensemble reweighting as previously described^39^. Fitted spectra were manually inspected and overlapped cross-peaks were excluded from further analysis. Sample integrity was assessed by interleaved ¹H–¹⁵N SORDID diffusion measurements, as previously described^35,42^.

Amide ¹H and ¹⁵N chemical shifts were obtained from 2D ¹H–¹⁵N SOFAST-HMQC experiments^78^ with a 50 ms acquisition time and 50 ms inter-scan delay. C′ shifts from BEST-HNCO experiments recorded at 700 MHz (acquisition ∼50 ms, inter-scan delay 200 ms), as previously described^39^.

### Molecular Dynamics Simulations

All MD simulations were performed using GROMACS (version 2021)^79^. Unbiased MD simulations of the isolated, unfolded state of FLN5 C747V were performed with the CHARMM36m force field with an increased protein-water Lennard Jones interaction strength (C36m+W)^80^ as previously described for isolated FLN5 A3A3^39^. We performed 15x 2 µs (30 µs total) of production simulations. Unbiased MD simulations of the WT nascent chain were performed using the C36m+W parameters, as previously described^39^. The ribosome was modelled from PDB 4YBB and truncated to retain atoms near the exit tunnel and exposed surface. The FLN6 linker and SecM were modelled from a cryo-EM map of FLN5+47 RNC, and the FLN5 domain was built from PDB 1QFH. Simulations of +31 WT RNC (C747V) included ∼1 million water molecules and 706 Mg²⁺ ions, resulting in a system size of ∼3.16 million atoms.

After initial minimisation, we performed NVT and NPT equilibration with position restraints on all heavy atoms, followed by restrained equilibration with only ribosome atoms position-restrained. A 2 fs timestep and LINCS constraints were used. Short-range electrostatics were calculated up to a cut-off of 1.2 nm, and the long-range component was computed using PME^81^, and van der Waals interactions were treated with a 1.2 nm cutoff and a switching function at 1.0 nm. Ten production simulations of 1.5 µs each (15 µs total) were run from independently equilibrated starting structures.

### PRE Calculations and Reweighting

PREs were back-calculated from MD ensembles as previously described^39^, incorporating τ_c,eff_ values that account for restricted tumbling of the nascent chain near the ribosome. The order parameter S^²^_NC_ was computed for each conformer to estimate τ_c,eff_ as a weighted average between isolated and ribosome-anchored τ_r_ values. PRE rates (Γ₂) were calculated using a rotamer library for MTSL and averaged over each ensemble. PRE intensity ratios (I_para_/I_dia_) were calculated as previously described^39^.

Four classes of PRE transverse relaxation rates (Γ_2_) were used for Bayesian reweighting of the wild-type FLN5+31 unfolded state structural ensemble. Cross-peaks with intensity ratios between 0.2-0.8 were restrained as the calculated Γ_2_ with propagated upper and lower bound relative errors derived as standard deviation of the spectral noise. Cross-peaks with intensity ratios between 0.1-0.2 and 0.8-0.9 were restrained in a similar manner with an addition of 5% propagated lower and upper bound errors. Cross-peaks with an intensity ratio >0.9 were restrained only with an upper bound of 3.39 s^-1^; cross-peaks with intensity ratios <0.1 were restrained only with a lower bound of 91.06 s^-1^.

Bayesian ensemble reweighting was performed with “Bayesian inference of ensembles” (BioEn)^82,83^ as previously described^39^, using experimental Γ₂ values as restraints. For RNCs, τ_c,eff_ and S^²^_NC_ were updated iteratively over 20 rounds to account for conformer-dependent effects. For ribosomal sites, the R_2_ rotamer distribution (better fitting to known structures) was used to improve agreement with intermolecular PREs.

### Structural Analysis

R_g_, SASA, contact maps, and secondary structure content were calculated using MDAnalysis (v2.9.0)^84^, MDTraj (v1.10.0)^85^, and GROMACS (version 2021)^79^. For the intrachain contact analysis, we defined a contact between two Cα atoms if the distances were less than 10Å, and contacts between closest heavy atoms and closest sidechain heavy atoms, with distances of less than 6Å. Native contacts were defined based on the FLN5 crystal structure as previously described^39^. Average contact probabilities were calculated as the mean Cα-Cα contact probability for residue pairs separated by at least 2 residues (total contacts) and 10 residues (native contacts), with errors representing the standard error of the mean. Secondary structure populations were calculated using DSSP^86^ implemented in MDTraj. Structural clustering was performed using the GROMOS algorithm, implemented in GROMACS, on unaligned nascent chain coordinates with a 2.2 nm RMSD cut-off. The ribosome–NC contacts were identified using a 12.5 Å threshold between NC Cα atoms and ribosome atoms: Cα atoms (protein) and P/N3/C4’ atoms (rRNA), as described previously^39^. Circos plots that visualise these interactions were generated as previously described^45,87^; the extent of ribosome-NC interactions was defined as % of frames in which a given contact was observed (after reweighting) and encoded in the greyscale intensity of the connecting lines. For better visualisation, we excluded contacts occurring in ≤1% of frames and adjusted the greyscale such that the darkest colour corresponds to 80%. For each NC residue, we calculated time series of native, non-native, and ribosome contacts involving that residue and visualised them as 2D correlation plots. Dimensionality reduction of ensembles was performed using the energy landscape visualisation method (ELViM)^54^ where only residues within the FLN5 domain of the nascent chain were used in this analysis for WT FLN5 and A3A3 ensembles. The 2D lower-dimension projections of each ensemble was used to map frames assigned to sub-ensembles (anchored, expanded and compact). We calculated 2D free energy surface of the ELViM projections of sub-ensemble indices on the density of indices in the ELViM projection^54^ and frames were extracted from the surface contours for downstream analysis of the most densely populated minima (<2 kJ/mol).

## Data availability

All NMR data are available as source data with the figures. The structural ensembles of the unfolded states will be deposited on Zenodo upon publication.

## Code availability

Example python scripts used to calculate PRE-NMR data from the ensembles and to refine the ensembles by reweighting are available on Github (https://github.com/julian-streit/PREreweighting).

## Acknowledgements

We acknowledge use of the UCL Biomolecular NMR Centre. This work was supported by a Wellcome Trust Investigator Award (to J.C., 206409/Z/17/Z) and by the Francis Crick Institute through provision of access to the MRC Biomedical NMR Centre. The Francis Crick Institute receives its core funding from Cancer Research UK (FC001029), the UK Medical Research Council (FC001029) and the Wellcome Trust (FC001029). This project made use of time on HPC resources on Archer2 (ARCHER2 UK National Supercomputing service, https://www.archer2.ac.uk) granted via the UK High-End Computing Consortium for Biomolecular Simulation, HECBioSim (http://hecbiosim.ac.uk), supported by EPSRC (grant no. EP/R029407/1 and EP/X035603/1). We also acknowledge the EuroHPC Joint Undertaking for awarding this project access to the EuroHPC supercomputer LUMI, hosted by CSC (Finland) and the LUMI consortium through a EuroHPC Regular Access call and the Baskerville Tier 2 HPC service (https://www.baskerville.ac.uk/). Baskerville was funded by the EPSRC and UKRI through the World Class Labs scheme (EP/T022221/1) and the Digital Research Infrastructure programme (EP/W032244/1) and is operated by Advanced Research Computing at the University of Birmingham. We additionally acknowledge the use of the UCL Myriad and Kathleen High Performance Computing Facility (Myriad@UCL and Kathleen@UCL), and associated support services, in the completion of this work.

## Author contributions

I.V.B., J.O.S., A.M.E.C. and J.C. designed and conceptualised the project. I.V.B., J.O.S. and A.M.E.C. produced the samples and performed the NMR experiments. I.V.B. analysed the NMR data. J.O.S. and T.W. performed and analysed the MD simulations. C.H. performed the ELViM analyses. S.H.S.C acquired and analysed the secondary chemical shifts used to validate the wild-type unfolded state ensemble on the ribosome. J.O.S., T.W. and J.C. sourced the computational resources and acquired the funding. I.V.B. and J.C. supervised the project. I.V.B., J.O.S., and J.C. administered the project. I.V.B. wrote the original draft. I.V.B. and J.C. edited the manuscript, and all authors reviewed the manuscript.

